# Novel newt regeneration genes regulate Wingless signaling to restore patterning in *Drosophila* eye

**DOI:** 10.1101/2021.02.28.433269

**Authors:** Abijeet Singh Mehta, Prajakta Deshpande, Anuradha Venkatakrishnan Chimata, Panagiotis A. Tsonis, Amit Singh

## Abstract

A fundamental process of regeneration, which varies among animals, recruits conserved signaling pathways to restore missing parts. Only a few animals like newts can repeatedly regenerate lost body parts throughout their lifespan that can be attributed to strategic regulation of conserved signaling pathways by newt’s regeneration tool-kit genes. Here we report use of genetically tractable *Drosophila* eye model to demonstrate the regeneration potential of a group of unique protein(s) from newt (*Notophthalmus viridescens*), which when ectopically expressed can significantly rescue missing photoreceptor cells in a *Drosophila* eye mutant. These newt proteins with signal peptides motifs exhibit non-cell-autonomous rescue properties and their regeneration potential even extends into later stages of fly development. Ectopic expression of these newt genes can rescue eye mutant phenotype by promoting cell proliferation and blocking cell death. These novel newt genes downregulate evolutionarily conserved Wingless (Wg)/Wnt signaling pathway to promote rescue. Modulation of Wg/Wnt signaling levels by using antagonists or agonists of Wg/Wnt signaling pathway in eye mutant background where newt gene(s) is ectopically expressed suggests that Wg signaling acts downstream of newt genes. Our data highlights the regeneration potential of novel newt proteins that regulate conserved pathways to trigger a robust regeneration response in *Drosophila* model with weak regeneration capability.

## Introduction

A urodele amphibian, newts belongs to *Salamandridae* family (Weisrock et al., 2006). The newts are the only four-legged vertebrates that exhibits a remarkable capability to regenerate tissues like limbs, tail, heart, lens, spinal cord, brain, and retina throughout its life time (Mehta and Singh, 2019a; Sanchez Alvarado and Tsonis, 2006). This regeneration potential/ ability of the newt is due to their ability to reprogram and dedifferentiate the terminally differentiated cells to trigger regeneration response (Mehta and Singh, 2019a). Such an exceptional regeneration ability of newts has been attributed to unique gene(s) that may have evolved to form the regeneration tool box (Bryant et al., 2017; Casco-Robles et al., 2018; Elewa et al., 2017; Evans et al., 2018; Keinath et al., 2015; Kumar et al., 2007; Matsunami et al., 2019; Mehta and Singh, 2019a; Nowoshilow et al., 2018; Sanor et al., 2020; Smith et al., 2009; Smith et al., 2019). Earlier, a newt gene Prod1, which encodes a transmembrane receptor, was found to be critical for maintaining proximo-distal identity (pattern memory) during newt limb regeneration (da Silva et al., 2002; Echeverri and Tanaka, 2005; Kumar et al., 2007; Mehta and Singh, 2019a). Homologs of Prod1 are yet to be identified outside of salamanders (Garza-Garcia et al., 2009). Similarly, the transcript of a novel newt gene *newtic1* was found to be significantly enriched in a subset of erythrocytes which formed an aggregate structure called erythrocyte clump. These *newtic* expressing erythrocytes promote limb regeneration (Casco-Robles et al., 2018). Surprisingly, our understanding of underlying molecular genetic mechanism(s) responsible for promoting lifelong regeneration of missing structures and/or rescue of pattern defects in newts is far from complete. Some strategies to determine their regeneration potential are to introduce these unique genes in genetically tractable model(s) that lack regeneration potential exhibited by newts such as mammals, *Drosophila* etc. Since the genetic machinery is conserved across the species, the *Drosophila* model has been successfully used for such cross species studies to model human disease and determine their underlying molecular genetic mechanism(s) (Hughes et al., 2012; Sarkar et al., 2018; Sarkar et al., 2016; Singh and Irvine, 2012; Tare et al., 2011).

Previously, a novel family of five protein members was identified in a comprehensive transcriptomic analysis from *Notophthalmus viridescens* (red-spotted newt). These proteins are expressed in adult tissues and are modulated during lens regeneration (Looso et al., 2013). The three-dimensional structure models of these novel newt proteins by *ab initio* methods suggested that these proteins could act as ion transporters and/or involved in signaling (Mehta et al., 2019). These unique newt genes were introduced in genetically tractable *Drosophila melanogaster* (fruit fly) model by transgenic approaches to test their regeneration potential. Such transgenic approaches were not feasible in *N*. *viridescens*, therefore, *Drosophila* served as a suitable system to address these queries. The transcriptomic analysis was performed in the transgenic *Drosophila* that are expressing newt genes. Based on these studies, graded expression of 2775 transcripts was reported in transgenic fly where one of the five newly identified newt genes was ectopically expressed using the GAL4/UAS binary target system (Brand and Perrimon, 1993; Mehta et al., 2019). Among these 2775 transcripts, genes involved in the fundamental developmental processes like cell cycle, apoptosis, and immune response were highly enriched suggesting that these foreign genes from newt were able to modulate the expression of *Drosophila* gene(s) (Mehta et al., 2019; Mehta and Singh, 2019b).

*Drosophila*, a hemimetabolous insect, serves as a versatile model organism to study development and disease due to presence of fully sequenced and less redundant genome, presence of homologs or orthologs for human disease related genes, a shorter life cycle and economy to maintain fly cultures (Bier, 2005; Lenz et al., 2013; Singh and Irvine, 2012). *Drosophila* eye has been extensively used to understand the fundamental process of development and model neurodegenerative disorders (Gogia et al., 2020a; Kumar, 2018; Sarkar et al., 2018; Sarkar et al., 2016; Singh and Irvine, 2012; Tare et al., 2011; Yeates et al., 2019). The rich repository of genetic tools available in fly model, and the fact that eye is dispensable for survival, makes *Drosophila* eye suitable for screening genes from other animals that can promote growth and regeneration. The compound eye of the adult fly develops from a monolayer epithelium called the eye-antennal imaginal disc, which develops from an embryonic ectoderm (Cohen, 1993; Held, 2002). The imaginal disc, a sac-like structure present inside the larva, comprise of two different layers: the peripodial membrane (PM) and the disc proper (DP) (Kumar, 2020b). The DP develops into retina whereas the PM of the eye-antennal imaginal disc contributes to the adult head structures (Haynie and Bryant, 1986; Kumar, 2020b; Milner et al., 1983). In third instar larval eye imaginal disc, a wave of synchronous retinal differentiation moves anteriorly from the posterior margin of the eye disc, and is referred to as Morphogenetic Furrow (MF) (Kumar, 2013, 2020a; Ready et al., 1976). The progression of MF transforms the undifferentiated cells into differentiated photoreceptor cells. The adult compound eye comprises of 800 unit eyes or ommatidia (Kumar, 2013, 2018, 2020a; Ready et al., 1976). The ommatidial cells in the compound eye are grouped together into two chiral forms, which are arranged in mirror image symmetry along the Dorso-Ventral (DV) midline called the equator. Ventral domain is the default state of the entire early eye imaginal primordium (Singh and Choi, 2003), and onset of the expression of the dorsal selector gene *pannier* (*pnr*), establishes the dorso-ventral (DV) lineage in the eye (Maurel-Zaffran and Treisman, 2000; Singh and Choi, 2003; Tare et al., 2013). It has been reported that *Lobe* (*L*) is a cell survival gene required for ventral eye development and growth, and acts upstream to the Wg signaling pathway (Singh et al., 2006).

The *wg* gene, a *Drosophila* homolog of Wnt, serves as a ligand, and encodes a secreted signaling protein that acts as a morphogen (Baker, 1987; Bejsovec, 2013; Swarup and Verheyen, 2012). The transcriptional effector of the Wg/β-catenin signaling is *Drosophila* T-cell Factor (dTCF), which upon activation by Armadillo (Arm, human homolog β-catenin) leads to the transcription of Wg target genes. In the absence of ligand Wg, a destruction complex comprised of Adenomatous polyposis coli (Apc), and Axin degrades Arm and thus prevents it to form a complex with dTCF. The pathway is activated by binding of Wg ligand to its co-receptors Frizzled (Fz) and Arrow (Arr). Upon activation, Fz binds to Dishevelled (Dsh) and Arr, which binds to Axin and thus inactivating the destruction complex. This results in the translocation of Arm (β-catenin) from the cytosol to the nucleus where Arm binds to dTCF and activates transcription of Wg target genes (Bejsovec, 2006). In the eye-antennal disc, Wg acts as a negative regulator of the eye development (Ma and Moses, 1995; Treisman and Rubin, 1995). In early third instar eye-antennal imaginal disc, *wg* expresses on the ventral margin. Ectopic expression of Wg triggers cell death of photoreceptor cells causing eye defects (Baker, 1987; Cavodeassi et al., 1999; Singh, 2012; Singh et al., 2006). Previously it has been shown that Wg pathway was upregulated in the developing eye of *L^2^* background. This perturbation in Wg pathway causes loss of ventral eye in 100% of flies of *L^2^*/+ background (Gogia et al., 2020a; Singh et al., 2006; Singh et al., 2011). Therefore, timing and level of *wg* expression in this region is crucial for formation of a normally patterned eye. Here, we demonstrate using a unique gene pool of newt genes (5 newly identified newt genes) to rescue *Drosophila L^2^* mutant pattern defect.

Misexpression of these novel newt gene in *Drosophila* downregulate Wg in *L^2^* mutant background, resulting in significant restoration of missing photoreceptor cells in the developing *Drosophila* eye. Ectopic expression of these genes restore eye mutant phenotypes by promoting cell proliferation, and downregulating cell death. Furthermore, these eye mutant phenotype rescues by newt genes are non-cell-autonomous and are all along the developmental stages *viz.,* early as well as later time points of development.

## Results

We employed *Drosophila* eye model to test the regeneration potential of these five unique newt genes, *candidate 1* (*C1), candidate 2 (C2), candidate 3 (C3), candidate 4 (C4),* and *candidate 5 (C5)*. We isolated the total RNA from newt tail, generated cDNA to amplify these newt genes, and generated transgenic flies for these five genes (Figure 1A). We employed GAL4/UAS system (Brand and Perrimon, 1993) to allow targeted misexpression of these newt transgenes in the fly tissues (Figure 1B). To confirm if targeted misexpression approach can generate newt proteins in *Drosophila* (Brand and Perrimon, 1993), we tested localization of the fusion protein. As there are no antibodies available against these novel newt proteins, so we used V5 epitope tagged transgenes to generate transgenic flies (Wang et al., 2019). The V5 tag sequence of about 42 bp long is fused towards the 3’ end of the 501 bp long open reading frame (ORF). We tested all these transgenes using the antibody against V5 tag. Here we present the expression of one of the transgenes *viz.*, C4 using V5 tag (Figure 1–figure Supplement 1). Next, to detect if misexpression of these newt genes in *Drosophila* can generate any developmental defects, we used *tubulin* – GAL4 (*tub* –GAL4) to ubiquitously misexpress newt genes in flies in the wild-type background (Figure 1B), and found no defects.

**Figure 1.**
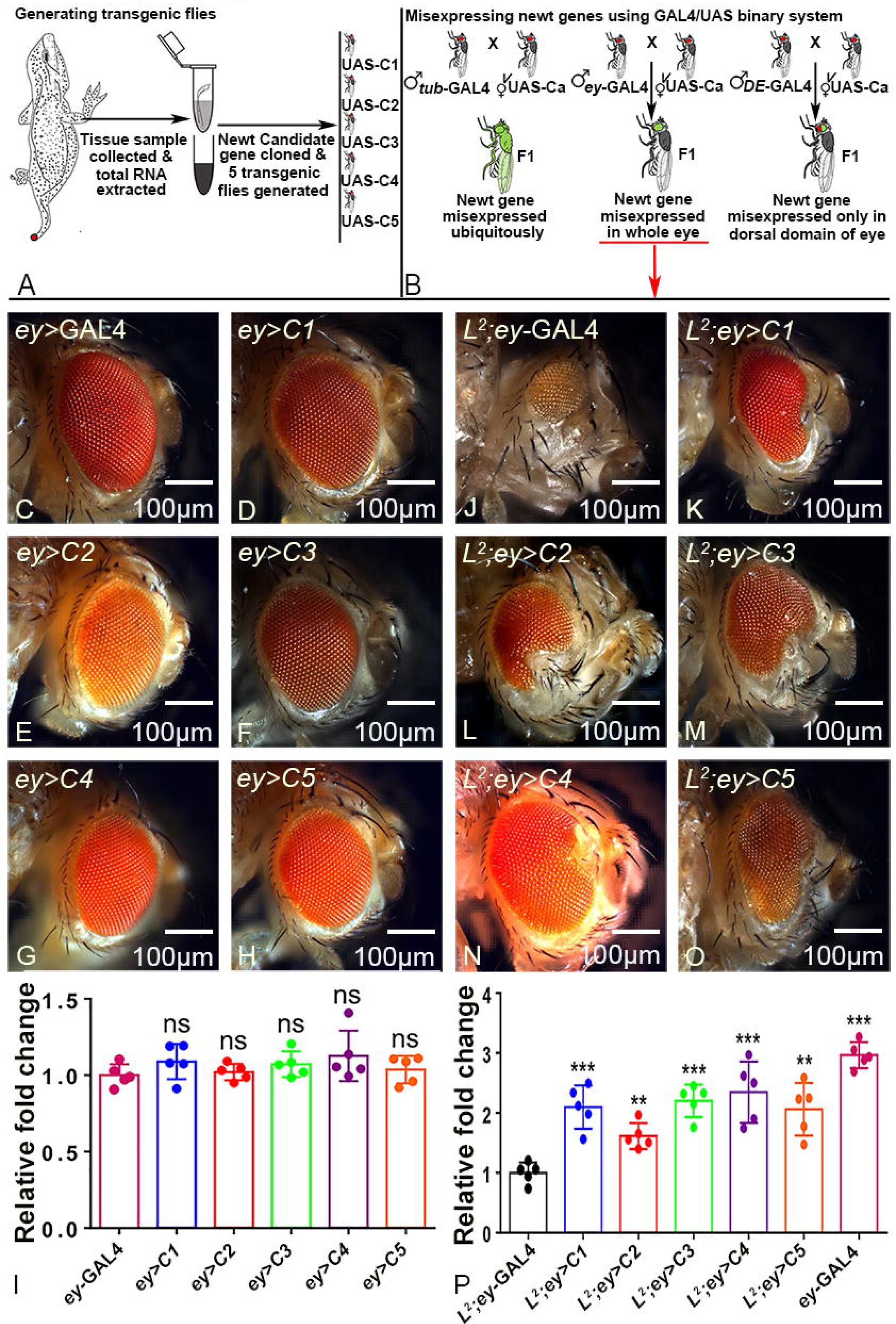
Novel newt genes rescues *L^2^* mutant’s loss-of-ventral eye phenotype. **(A, B)** Schematic representation of generating transgenic *Drosophila* carrying novel newt (*Notophthalmus viridescens*) candidate genes (*C1, C2, C3, C4,* and *C5*). Bright field image of a **(C)** wild type adult eye as positive control (*ey-*GAL4), (D-H) adult eye phenotype when *ey*-Gal4 drives expression of newt transgenes **(D)** UAS-*C1 (ey>C1)*, **(E)** UAS-*C2* (*ey>C2*), **(F)** UAS-*C3 (ey>C3)*, **(G)** UAS-*C4 (ey>C4)*, and **(H)** UAS-*C5 (ey>C5)*. **(I)** The bar graph shows there is no significant (ns) change in eye size between control and where ey-GAL4 drives expression of newt transgenes (*ey>C1-C5*). Bright field image of a **(J)** *L^2^* mutant (*L^2^; ey-*GAL4) adult eye where there is preferential loss-of-ventral eye phenotype, **(K-O)** *L^2^* mutant (*L^2^; ey-*GAL4) background where newt transgene **(K)** UAS-*C1 (L^2^; ey >C1)*, **(L)** UAS-*C2* (*L^2^; ey >C2*), **(M)** UAS-*C3 (L^2^; ey >C3)*, **(N)** UAS-*C4 (L^2^;ey>C4)*, and **(O)** UAS-*C5 (L^2^;ey>C5)* are expressed in the developing eye using ey-GAL4. **(P)** *L^2^* mutant adult eye is half the adult eye size of the *ey-*GAL4 which serve as positive control. The bar graph clearly displays the significant increase in the adult eye size of *L^2^-*mutant background where unique newt genes were misexpressed compared to the adult eye size of the *L^2^-*mutant only (*L^2^; ey-*GAL4). Eye size was measured using ImageJ software tools (http://rsb.info.nih.gov/ij/). Sample size used for statistical analysis was five (n=5). Statistical analysis was performed using student’s t-test for independent samples. Statistical significance was determined with 95% confidence (p<0.05). Error bars show standard deviation, and symbols above the error bar signify as: ns is non-significant p value, * signifies p value <0.05, ** signifies p value <0.005, *** signifies p value <0.0005 respectively. In this study *ey* is used as a driver to misexpress *newt genes* in the developing *Drosophila* eye. All the images are displayed in same polarity as dorsal domain-towards top, and ventral domain-towards bottom. Scale bar= 100 μm.

### Newt genes can rescue *Drosophila* eye mutants

For our experiments, we used the *Drosophila* eye model as it is not required for the viability of the fly and it is easy to score the phenotypes. We used *ey*-GAL4 driver (Hazelett et al., 1998) to misexpress these newt genes in the developing eye imaginal disc, which develops into an adult eye. The phenotypes of UAS-transgene alone were tested to rule out any contribution(s) in phenotype due to insertion of the transgene. In comparison to the control, *ey*-GAL4 (Figure 1C), misexpression of these transgenes *ey*-GAL4>UAS-*C1* (*ey*>*C1*, Figure 1D), *ey*-GAL4>UAS-*C2 (ey*>*C2*, Figure 1E), *ey*-GAL4>UAS-*C3* (*ey*>*C3*, Figure 1F), *ey*-GAL4>UAS-*C4* (*ey*>*C4*, Figure 1G), and *ey*-GAL4>UAS-*C5* (*ey*>*C5*, Figure 1H), did not affect the eye size (Figure 1I). To investigate the regeneration potential of these genes, we used eye mutant(s) such as *L* mutant which exhibits selective loss of ventral half of the eye (Figure 1J). This mutant was selected as the penetrance of the phenotype of loss-of-ventral eye is 100%. Interestingly, targeted misexpression of the newt gene(s) in *L^2^*; *ey*-GAL4 background [(*L^2^*; *ey*>*C1*; Figure. 1K), (*L^2^; ey>C2;* Figure 1L), (*L^2^; ey>C3;* Figure 1M), (*L^2^; ey>C4*; Figure 1N), (*L^2^; ey>C5*; Figure 1O)] exhibit significant rescue of the loss-of-ventral eye phenotype (Figure 1P). For each cross, three independent cultures were established and two hundred flies for each set (triplicate, 600 flies) were counted to determine phenotype frequency. The rescue frequency was about 40.83% in C1, 37.7% in C2, 49% in C3, 58.2% in C4, and 38.7% in C5 (Figure 1–figure Supplement 2). Note that the strongest rescues were observed with C4 transgene (Figure 1P; F1–S2; and Table S1).

Since the eye mutant represent developmental defects in the genetic machinery involved in early eye development, it raises a question whether the regeneration potential of these newt genes is restricted to the early time window or even later stages of eye development. The GMR-GAL4 driver was used to drive expression of transgenes in the differentiating retinal neurons of larval eye imaginal disc, which continues all along to the adult eye (Moses and Rubin, 1991). Gain-of-function of *head involution defective* (*hid*), an executioner caspase of cell death machinery, triggers cell death (Bergmann et al., 1998; Grether et al., 1995; Hay et al., 1995; White et al., 1994). Misexpression of *hid* using GMR-GAL4 (GMR>*hid*) results in a “No-eye” or highly reduced eye phenotype (Figure 1–figure Supplement 3B). However, misexpression of newt transgenes along with *hid* in the eye (GMR-GAL4>*hid*+*C4*) results in a significant rescue of the “No-eye” phenotype (Figure 1–figure Supplement 3C). Since these newt genes have signal peptides (Looso et al., 2013), it is possible that these genes may have non-autonomous effects.

### Ectopic expression of newt genes exhibits non-autonomous rescue

To determine if these genes have non-autonomous effect, we misexpressed these genes within a subset of retinal neuron population in the developing eye field of *L^2^* mutant and assay their regeneration effect. Misexpression of *C4* using *dorsal-eye -* GAL4 (*DE -* GAL4) (Figure 1B) directs its expression in the dorsal half of the developing eye (Morrison and Halder, 2010) (Figure 2A, E). The rationale of this experiment was to misexpress *C4* in the dorsal half of the eye and test its effect on an eye mutant which results in the loss of the ventral half of the developing eye (Figure 2). Misexpression of both GFP (reporter) and newt *C4* in the dorsal domain of the developing eye, which serves as a positive control (*+*/+*; DE>GFP+C4)* (Figure 2A, B, B′, B″), exhibits a near wild-type eye. As a negative control, we tested GFP expression alone in *L^2^*-mutant background (*L^2^*/+*; DE>GFP)* (Figure 2C, D, D′, D″). We found that GFP expression was restricted to the dorsal half as seen in the wild-type control (Figure 2B, B′, B″). Furthermore, we did not see any rescue of *L* mutant phenotype of ventral eye loss (marked by red dotted boundary, Figure 2B, B′, B″). Misexpression of both GFP+C4 in the dorsal half of the *L^2^* mutant eye (*L^2^*/+*; DE>GFP+C4)* exhibits a significant rescue of the loss-of-ventral eye phenotype as seen in the adult eye, and the eye imaginal disc (marked by white-dotted boundary, Figure 2E, F, F′, F″). Even though the targeted misexpression of C4 is restricted to the dorsal half of the developing eye, it was able to rescue the loss-of-ventral eye phenotype suggesting that newt genes can show non-cell-autonomous rescue of *Drosophila* eye mutants.

**Figure 2.**
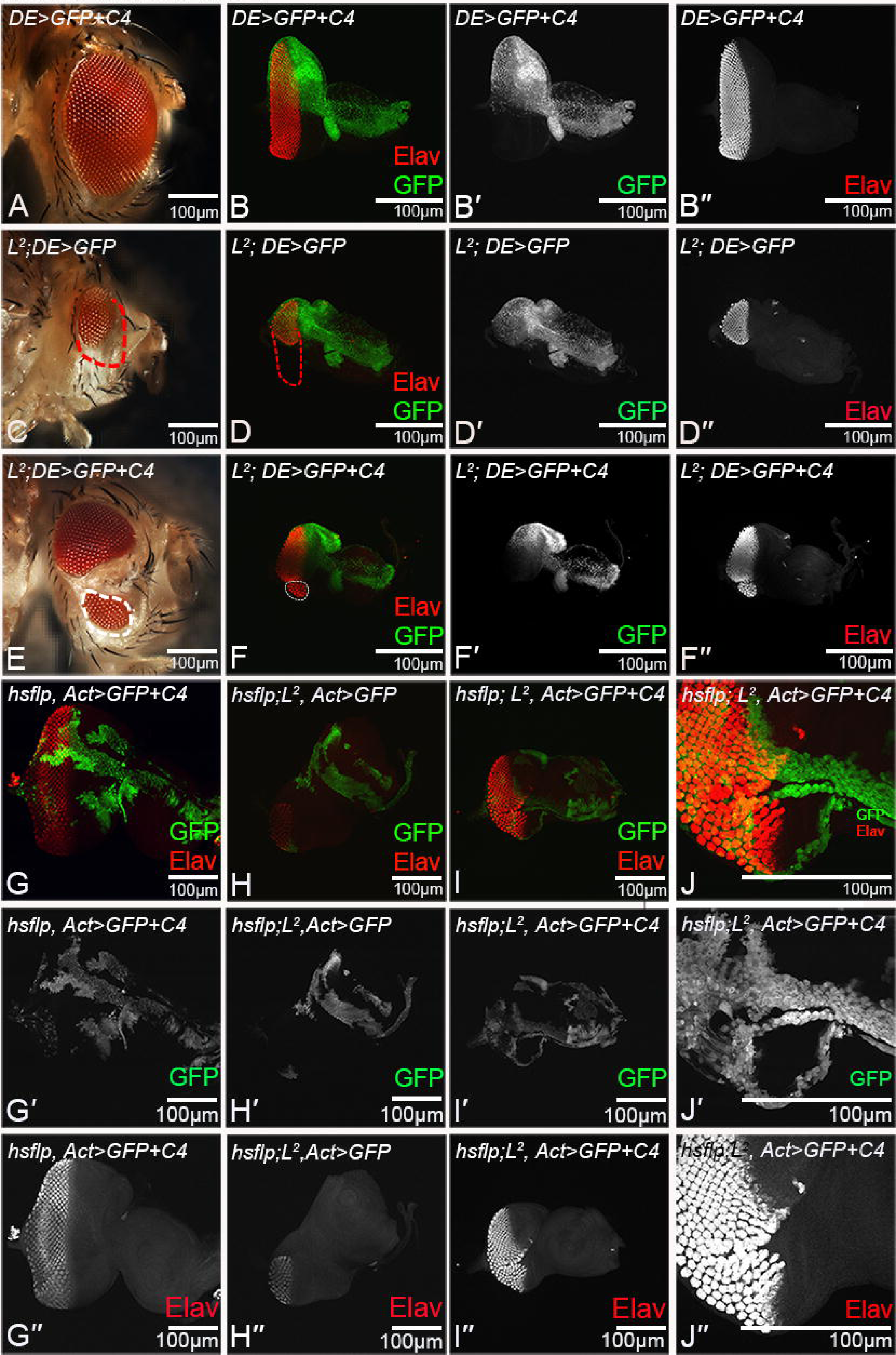
Misexpression of newt genes exhibits non-cell-autonomous rescue. **(A, B, B′, B″)** Misexpression of *C4* in the dorsal half using *DE-*GAL4 driver results in **(A)** Adult eye and **(B, B**′**, B**″) developing third instar larval eye imaginal disc (as positive control). Note that Green fluorescence protein (GFP) (green) reporter marks the DE-GAL4 driver domain and proneural marker Embryonic lethal abnormal visual system (Elav) (red) marks the retinal fate. **(B**′) is a single channel confocal image showing GFP expression, and **(B**″) is a single channel confocal image showing Elav expression. **(C, D, D**′**, D**″) *L^2^* mutant eye where there is no misexpression of newt genes (*L^2^*/+*; DE>*GFP) (as negative control). **(C)** Adult eye bright field image and **(D, D**′**, D**″) third instar eye disc of *L^2^* mutant (*L^2^*/+*; DE>*GFP) exhibiting loss-of-ventral eye phenotype and red dotted line marks the boundary of eye. GFP reporter (green) marks domain of DE-GAL4 expression domain in *L^2^* mutant background. **(E, F, F**′**, F**″) Misexpression of C4 in dorsal eye domain of *L^2^* mutant (*L^2^*/+*; DE>*GFP; C4) exhibits non-cell autonomous rescue of loss-of-ventral eye phenotype in **(E)** Adult eye and **(F**′**, F**′**, F**″) third instar eye disc. White dotted boundary marks the significantly rescued loss-of-ventral eye phenotype. **(G-J)** Genetic mosaic “Flp out” somatic clones of **(G,G**′**,G**″) newt gene C4 in the developing eye disc wild-type, **(H,H**′**,H**″) GFP in *L^2^* background (*L^2^*/+*; hsflp>*GFP, served as the negative control) and **(I,I**′**,I**″ **& J,J**′**,J**″) newt gene *C4* clones in *L^2^* mutant (*L^2^*/+*; hsflp>*GFP*;C4*) background. All clones are marked by GFP reporter (green). Note that misexpressing *C4* rescue missing photoreceptor cells in a non-cell-autonomous manner, as photoreceptor cells rescue marked by Elav (in red) extends outside the boundary of the clone marked by GFP (in green). **(J, J**′**, J**″) exhibits the magnified view of the *C4* misexpressing clone (green) in *L^2^*/+*; hsflp>*GFP*; C4* background. All the images are displayed in same polarity as dorsal domain-towards top, and ventral domain-towards bottom. Scale bar= 100 μm.

In order to rule out any domain specific restraint, or contribution of a *DE-*GAL4 background, we employed genetic mosaic approach. We generated gain-of-function clones using Flp out approach (Xu and Rubin, 1993), which result in clones of cells that misexpress high levels of the transgene and these clones are marked by the GFP reporter (Figure 2G,H,I,J). We found that the misexpression of these transgenes rescues the loss-of-ventral eye phenotypes, however these rescues extended beyond the boundary of the clone. This data validate our prior observation that these newt gene exhibit non-cell-autonomous rescue due to their secretory nature. Next, we wanted to investigate the mechanism by which these newt genes can rescue loss-of-ventral eye phenotype.

### Newt genes promote cell proliferation and downregulate cell death

To test the possibility of upregulation of cell proliferation resulting in the rescue of the missing tissue in the *L^2^* mutant background, we calculated relative fold change in mitotic index (MI) of the eye imaginal discs showing rescue of *L* mutant phenotype and compared it to the control *L^2^* mutant (*L^2^*; *ey*-GAL4) eye disc (Figure 3A-D). Mitotic index is the ratio between the number of proliferating cells to the rows of ommatidial cells (Baonza and Freeman, 2005; Vollmer et al., 2016; Yang et al., 2005). The proliferating cells are marked by anti-phospho-Histone H3 (PH3) marker (Figure 3A-C, A′-C′ and Figure 3–figure Supplement 1A-F, A′-F′). Phosphorylation of a highly conserved Serine residue (Ser-10) in the histone H3 tail is considered a crucial event for the onset of mitosis, and represents completion of cell cycle (Crosio et al., 2002; Kim et al., 2017). Our results show robust upregulation of about 5.79 ± 1.5 folds increase in the MI when *C4* is misexpressed in the *L^2^-*mutant background (*L^2^; ey*>*C4*) with respect to the *L^2^-* mutant (*L^2^*; *ey*-GAL4) only (Figure 3D). Similar phenotypes were observed with other transgenes (Figure 3–figure Supplement 1A-G) and the MI values were *C3*: 5.18 ± 1.10, *C2:* 4.6 ± 1.7, *C1*: 4.3± 1.4, and *C5*: 4.2 ± 0.9 respectively. These results show that the newt gene(s) can induce cell proliferation to restore the missing photoreceptor cells of *L^2^*-mutant eye disc and thus exhibit significant rescue of the loss-of-ventral eye phenotype as seen in the eye imaginal disc and the adult eye (shown in white dotted boundary, Figure 3B). The control where no newt gene is misexpressed, in the *L^2^* mutant background did not show any rescue of the loss-of-ventral eye phenotype due to lack of cell proliferation as evident from PH3 staining (shown in red dotted boundary, Figure 3A).

**Figure 3.**
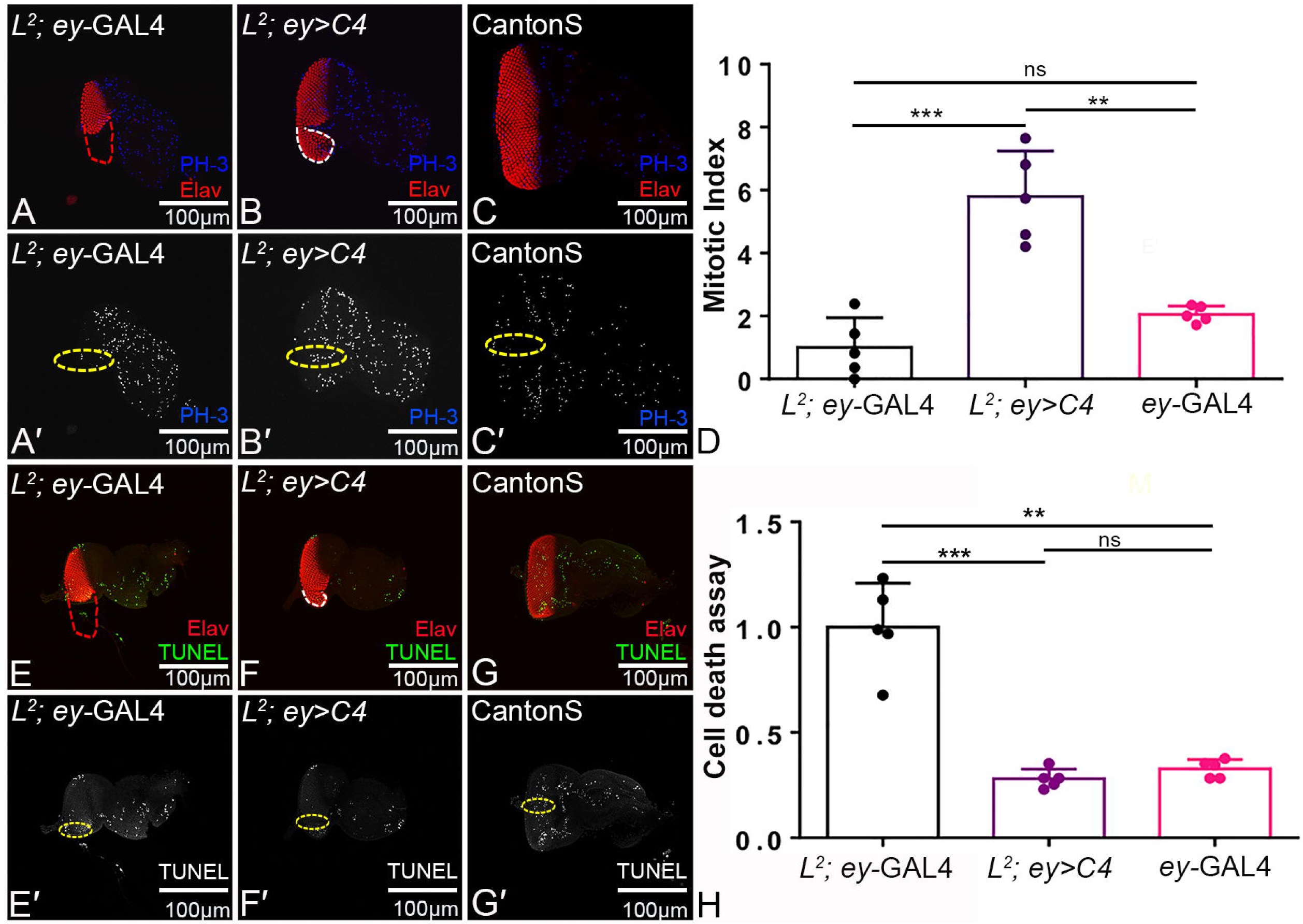
Newt genes promote cell proliferation and downregulate cell death to promote rescue of *L^2^* mutant phenotype of loss-of-ventral eye. **(A-D)** PH-3 staining (blue), as a marker to calculate cell proliferation and mitotic index, in (A) *L^2^* mutant eye disc (red dotted boundary marking eye boundary), (B) *L^2^* mutant eye disc where C4 gene is misexpressed (*L^2^*; *ey*>C4), and **(C)** Wild type (*ey*-GAL4) eye disc which serve as control. **(A**′**-C**′) single channel confocal images demonstrating PH-3 stained nuclei. **(B)** White dotted boundary represents the partially rescued loss-of-ventral eye phenotype. The zone of interest that was considered to count number of dividing cells is marked in yellow dotted boundary. **(D)** Bar graph showing relative increase in the mitotic index (MI) when newt genes are misexpressed in *L^2^* mutant background. MI is calculated as ratio of cells dividing (stained by PH-3 in blue) to total rows of ommatidial cells (stained by Elav in red). There is no significant change in MI between the *L*^2^ mutant, and wild-type (*ey*-GAL4) eye disc, but comparatively there is robust increase in the MI in*L^2^*; *ey*>C4 background. **(E-H)** Terminal deoxynucleotidyl transferase-mediated dUTP nick-end labeling (TUNEL) staining as a marker for cell death assay. TUNEL staining (shown in green) marks the dying nuclei in eye imaginal disc of (E) *L^2^* mutant, (F) *L^2^* mutant where C4 gene is misexpressed (*L^2^*; *ey*>*C4*), and **(G)** Wild type (*ey*-GAL4) eye disc. Red dotted boundary represents ventral-eye-loss area, and white dotted boundary represent the rescued loss-of-ventral eye region. **(E**′**-G**′) The tunnel positive cells are counted within the zone of interest marked within yellow dotted boundary as shown in single filter confocal images. **(H)** bar graph showing relative decrease in the cell death when newt gene is misexpressed under the *L^2^*-mutant background. The downregulation in cell death is comparable to the cell death happening in the normal wild type background. Statistical analysis was performed using student’s t-test for independent samples. Sample size used for the calculation was five in number (n=5). Statistical significance was determined with 95% confidence (p<0.05). All bar graphs show relative fold change in cell proliferation and cell death as average between 5 samples. Error bars show standard deviation, and symbols above the error bar ** signify p value <0.005, *** signify p value <0.0005, and ns signify p value > 0.05 respectively. All the images are displayed in same polarity as dorsa domain-towards top, and ventral domain-towards bottom. Scale bar= 100 μm.

During the newt lens regeneration both cell proliferation and apoptosis are seen (Tsonis et al., 2004), which warranted the need to look at the levels of cell death. It has also been shown that the loss-of-ventral-eye phenotype observed in *L^2^* mutation is due to induction of both caspase-dependent and caspase-independent cell death (Singh et al., 2006). Therefore, we tested if these newt genes can block or downregulate cell death to rescue *L^2^* mutant phenotype by using the TUNEL (Terminal deoxynucleotidyl transferase dUTP nick end labeling) assay (Figure 3E-G, and Figure 3–figure Supplement 1H-M). TUNEL assay is based on the detection of fragmented DNA ends, which are characteristic of apoptotic cells (Sarkissian et al., 2014; White et al., 2001). Our results clearly demonstrate strong decrease in the number of dying cells (0.28 ± 0.04 fold) when newt gene (*C4*) is misexpressed under the *L^2^* mutant background (*L^2^; ey*>*C4*) (Figure 3F, F′, H) relative to the *L^2^*-mutant strain in which newt gene is not misexpressed (*L^2^; ey*-GAL4) (Figure 3E, E′, H). The positive control (*ey*-GAL4) shows 0.32 ± 0.04 fold downregulation in the rate of cell death (Figure 3G, G′, H) relative to the *L^2^* mutant (*L^2^*; *ey*-GAL4). The value is significantly lower in comparison to the *L^2^* mutant *(L^2^*; *ey*-GAL4), but approximately equivalent to the *L^2^* mutant where newt gene is misexpressed (*L^2^; ey*>C4) (Figure 3H). This suggests that newt gene(s) can recue the *L^2^* mutant loss-of-ventral eye phenotype to wild-type by preventing excessive cell death of photoreceptor cells (marked in white dotted boundary, Figure 3 F, F′). We found similar results when other members of novel newt gene family were misexpressed (Figure 3–figure Supplement 1H-O). Downregulation in cell death is observed in C3: 0.36 ± 0.14, C2: 0.38 ± 0.09, C1: 0.39 ± 0.15, and C5: 0.45 ± 0.22 respectively (Figure 3–figure Supplement 1N). We also tested the role of cell death using Dcp1 (Death caspase-1) staining (Figure 3–figure Supplement 2). Antibody against cleaved Dcp1 marks the dying cells in *Drosophila* (Sarkissian et al., 2014). Dcp1 staining exhibits reduction in the number of dying cells when newt genes are misexpressed in *L^2^* mutant background (Figure 3–figure Supplement 2C-F), compared to when newt genes are not expressed in *L^2^* mutant background (Figure 3–figure Supplement 2B). These results suggest that newt genes upregulate cell proliferation and downregulate cell death to significantly rescue the *L^2^* mutant phenotype of loss-of-ventral eye.

We further validated the role of cell death by blocking caspase dependent cell death using baculovirus p35 misexpression (Hay et al., 1994). Even though misexpression of *p35* in *L^2^* mutant background (*L^2^*/+*; ey>p35*) exhibits rescue of loss-of-ventral eye phenotype, the phenotype strength (relative eye size ratio of 1.42 ± 0.29 and rescue frequency of 22.7%) of *L^2^*/+*; ey>p35* was significantly weaker than phenotype strength (relative eye size ratio = 2.34 ± 0.51 and rescue frequency of 58.2%) of *L^2^*/+*; ey>C4* (Figure 3–figure Supplement 3C, D, F, G). This data suggests that the newt genes upregulate cell proliferation and block cell death as well, but cell proliferation may be playing significant role in rescuing the missing tissues in *the L^2^* mutant eye. It is therefore important to understand what signal is triggered by these newt genes to promote the rescue.

### Newt genes downregulate *wg* expression in developing eye

We screened for the signaling pathway(s) that are involved in the rescue of *L^2^* mutant loss-of-ventral eye phenotype. We identified members of evolutionarily conserved Wg/Wnt signaling pathway in our high throughput screen (RNA sequencing) where newt genes were misexpressed in fly tissue (Figure 4–figure Supplement 1). It is reported that *L^2^* mutant phenotype can be rescued by downregulating *wg* (Singh et al., 2006). To determine the involvement of *wg* in promoting rescue of *L^2^* mutant phenotype by unique newt genes, we examined expression levels of Wg in larval eye-antennal imaginal disc (Figure 4A-C, A′-C′, D). Wg expression is seen at antero-lateral margins of the developing eye imaginal disc (Baker, 1987; Cavodeassi et al., 1999; Singh et al., 2006). In *L^2^* mutant robust ectopic expression of Wg is seen at the ventral eye margins (Figure 4B, B) (Singh et al., 2006; Singh et al., 2011). Misexpression of newt gene (*C4*) in *L^2^* mutant background downregulates the Wg expression at the ventral margin of eye-antennal imaginal disc (Figure 4C, C′), as evident from the signal intensity plots (integrative density) in various backgrounds (Figure 4D). The semi-quantitative western blot, showed that Wg levels increased 3 folds in *L^2^* mutant as compared to *ey-GAL4* control whereas misexpressing *C4* in the *L^2^* mutant background decreased the Wg expression by 1.6-fold as compared to *L^2^* mutant (*L^2^; ey*-GAL4) (Figure 4 E, F). Similar trends were seen with other novel newt genes (C1, C2, C3, and C5) when misexpressed in *L^2^* mutant background (Figure 4–figure Supplement 2C-F, C′-F′, G, G′). Thus, the newt genes can downregulate *wg* expression in *Drosophila L^2^* mutant to partially rescue ventral eye-loss-phenotype.

**Figure 4.**
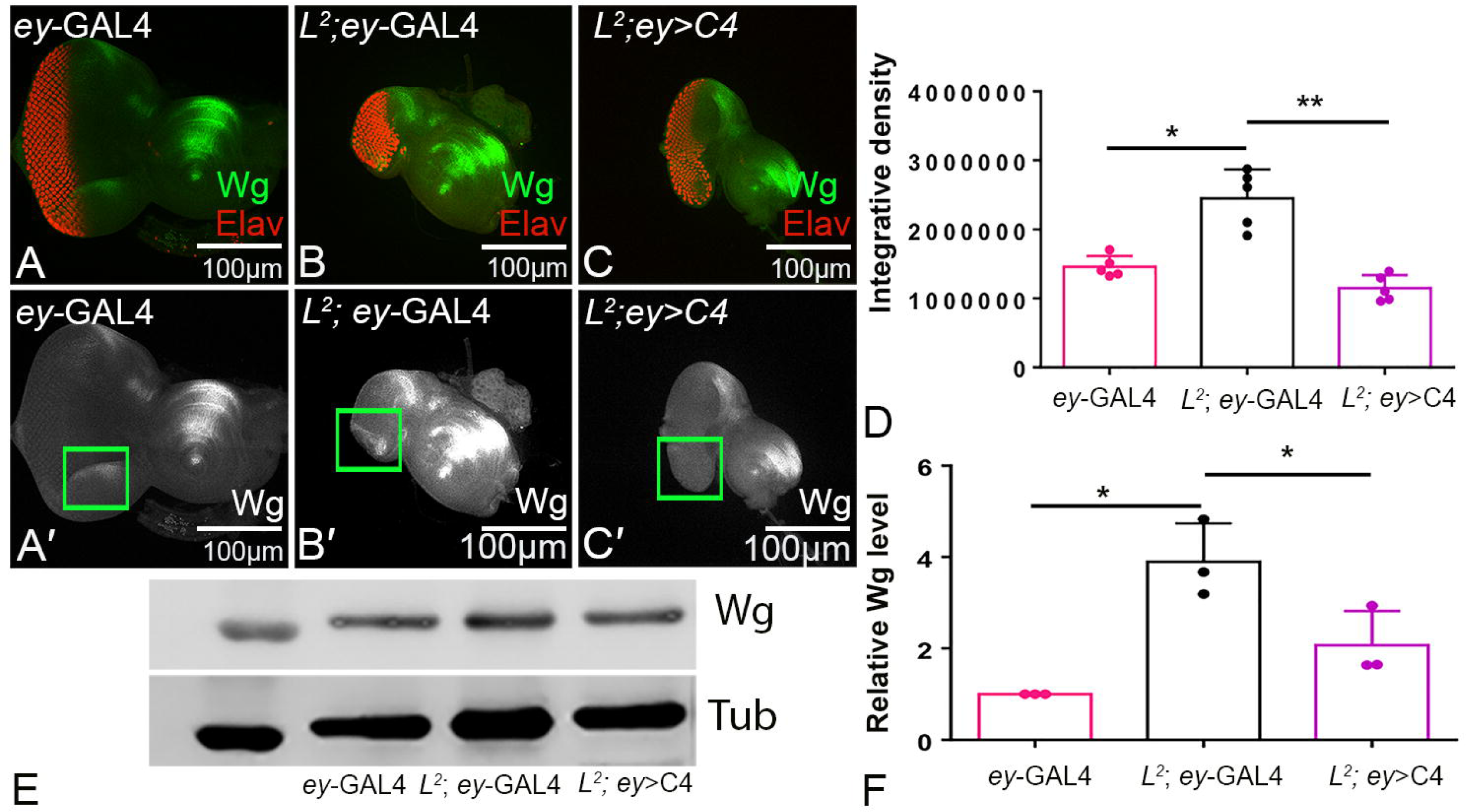
Newt genes downregulate Wg expression to rescue loss-of-ventral eye phenotype. **(A-D)** Immunohistochemistry showing Wg modulation. **(A-C)** Eye disc showing Wg staining in green, and Elav in red **(A**′**-C**′) are single channel confocal images showing Wg expression. The green window is the area of interest that is utilized to calculate **(D, D**′) integrative density (intensity plot). The bar graph represent the Wg intensity at the ventral margin (within the green box) in **(A, A**′) wild type fly (*ey*-GAL4**), (B, B**′) *L^2^*-mutant fly, **(C, C**′) Misexpression of newt gene (*C4*) in the *L^2^* mutation background. Note that *C4* misexpression downregulates Wg expression. **(E, F)** Western blot (WB) to semi quantitate modulation of Wg expression levels. **(E)** The blot shows the expression of Wg protein (C4-V5) in *Drosophila melanogaster.* The samples were loaded in the following sequence: Lane 1-*ey-*GAL4, Lane 2-*L^2^; ey-*GAL4, Lane 3-*L^2^; ey>C4.* Alpha tubulin is used as a loading control. Molecular weight of Wg is 46kD, and alpha tubulin is 55 kD. The samples were treated with anti-Wg antibody, and anti-α tubulin antibody. **(F)** Bar graph representing as a relative Wg level, is a measure of signal intensity of the bands, which clearly demonstrates that newt genes when misexpressed under the *L^2^*-mutant background downregulates Wg levels. Statistical analysis was performed using student’s t-test for independent samples. Statistical significance was determined with 95% confidence (p<0.05). All bar graphs show integrative density as a scale to measure Wg intensity for each sample represented as the average between 5. Error bars show standard deviation, and symbols above the error bar * signify p value <0.05, and ** signify p value <0.005 respectively. All the images are displayed in same polarity as dorsal domain-towards top, and ventral domain-towards bottom. Scale bar= 100 μm.

To further test our hypothesis, we misexpressed *wg* (the target identified in our screen) along with *C4* transgene in the *L^2^* mutant background. As control, we misexpressed *wg* in the developing eye (*ey>wg*), which result in reduced eye phenotype (Figure 5 A, B) (Lee and Treisman, 2001; Singh et al., 2012a). Misexpressing *C4* in the wild-type background where *wg* is ectopically induced (*+*/+*; ey>wg+C4*), rescues the reduced eye phenotype (Figure 5 C, D) at a frequency of 16.8% (Figure 5–figure Supplement 1 and Table S1). Misexpression of *wg* in the *L^2^*/+ mutants eye background (*L^2^*/+*; ey>wg*) completely eliminate the eye field as seen in the eye imaginal disc and the adult eye (Figure 5E, F) whereas, misexpressing *C4* along with *wg* in the *L^2^*/+ mutant eye disc (*L^2^*/+*; ey>wg; C4*) partially rescues the loss-of-ventral eye phenotype (Figure 5 G, H) in 21% flies (Figure 5–figure Supplement 1 and Table S1). Thus, newt genes can even downregulate Wg expression in the wild-type (*+*/+*; ey>wg+C4*; Figure 5 C, D) as well as *L^2^* (*L^2^*/+*; ey>wg; C4*; Figure 5 G, H) background(s) where *wg* was ectopically co-expressed in *Drosophila* eye to promote rescue of the loss-of-ventral eye phenotype.

**Figure 5.**
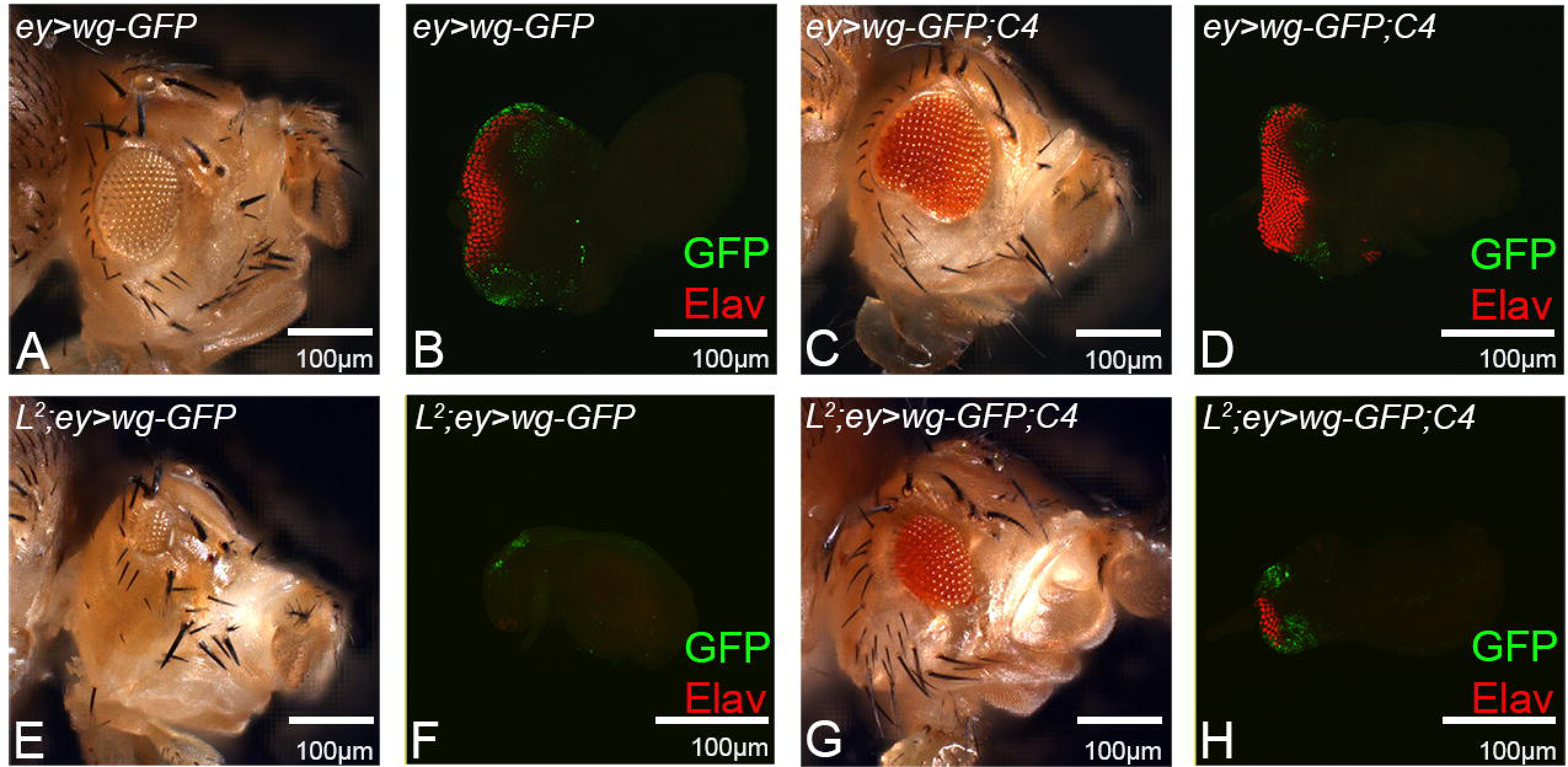
Newt gene regulates ectopically misexpressed Wg both under the wild type background, and under the *L*^2^-mutant background. **(A-H)** Wg is ectopically misexpressed in the *Drosophila* developing eye. **(A,B)** wild type background (*+*/+*; ey>wg-GFP*). **(C, D)** Newt gene and Wg are misexpressed under the wild type background (*+*/+*; ey>wg-GFP;C4*) showing increase in the eye size compared to (*+*/+*; ey>wg-GFP*). Comparatively represented as both **(A, C)** bright field adult eye picture, and (B, D) third instar eye-antennal imaginal disc confocal image. **(E, F)** Wg is misexpressed under the *L^2^*–mutant background (*L^2^*/+*; ey>wg-GFP*) that lack newt gene. *L^2^*–mutant small eye phenotype becomes more severe. **(G, H)** Newt gene and Wg are misexpressed under the *L^2^-mutant* background (*L^2^*/+*; ey>wg-GFP; C4*). There is a significant rescue in the eye-loss phenotype. Comparatively represented as both **(E, G)** bright field adult eye picture, and (F, H) third instar eye-antennal imaginal disc confocal image. The GFP as reporter is in green that signifies ectopic misexpression of Wg in the developing eye. Elav staining is shown in red. All the images are displayed in same polarity as dorsal domain-towards top, and ventral domain-towards the bottom. Scale bar= 100 μm.

### Newt genes negatively regulate Wg/Wnt signaling

To investigate if newt genes regulate expression of *wg* alone or they regulate the Wg signaling, we tested the antagonists, and agonists of the Wg-signaling pathway. The rationale was to modulate the levels of Wg signaling and sample its effect on the *L^2^* mutant phenotype as well as the *L^2^* mutant phenotype where *C4* is misexpressed (*L^2^*/+*; ey> C4)* (Figure 6). Increasing the levels of Arm (a vertebrate homolog of β-Catenin, an agonist of Wg signaling) in the wild-type eye (*ey>arm*) resulted in small eyes (Figure 6–figure Supplement 1A, B) and in the *L^2^*/+ background, (*L^2^*/+*; ey>arm)* it results in elimination of the entire eye field (Figure 6B, C). Misexpressing *C4* along with *arm* in the *L^2^*/+ mutant background (*L^2^*/+*; ey>arm; C4*) partially rescues the loss-of-ventral eye phenotype (Figure 6D, E) and the rescue frequency is 22.8% (Figure 6–figure Supplement 2A and Table S1). Similarly, misexpressing *C4* and *arm* in the wild-type background (*+*/+*; ey>arm; C4*) partially rescues the small eye phenotype defect caused by *ey>arm* phenotype (Figure 6–figure Supplement 1C, D), and the rescue frequency is 21% (Figure 6–figure Supplement 2A and Table S1). This data suggests that newt gene can modulate the Wg signaling pathway components to promote rescue of reduced eye phenotype both in *L*^2^-mutant as well as the wild-type background(s). However, the rescue frequency obtained by misexpressing newt gene in addition to ectopically misexpressing *arm* (*L^2^*/+*; ey>arm; C4*, 22.8%) and/or *wg* (*L^2^*/+*; ey>wg; C4*, 21%) (Figure 5–figure Supplement 1, Figure 6–figure Supplement 2A and Table S1) is lower than the rescue frequency obtained when only newt gene is misexpressed in the *L^2^-*mutant background (*L^2^*/+*; ey>arm; C4*, 58.2%). This could be due to severity of loss-of-ventral eye phenotype caused by the combined effect of *L^2^* mutation and gain-of-function of *wg*, or *arm*.

**Figure 6.**
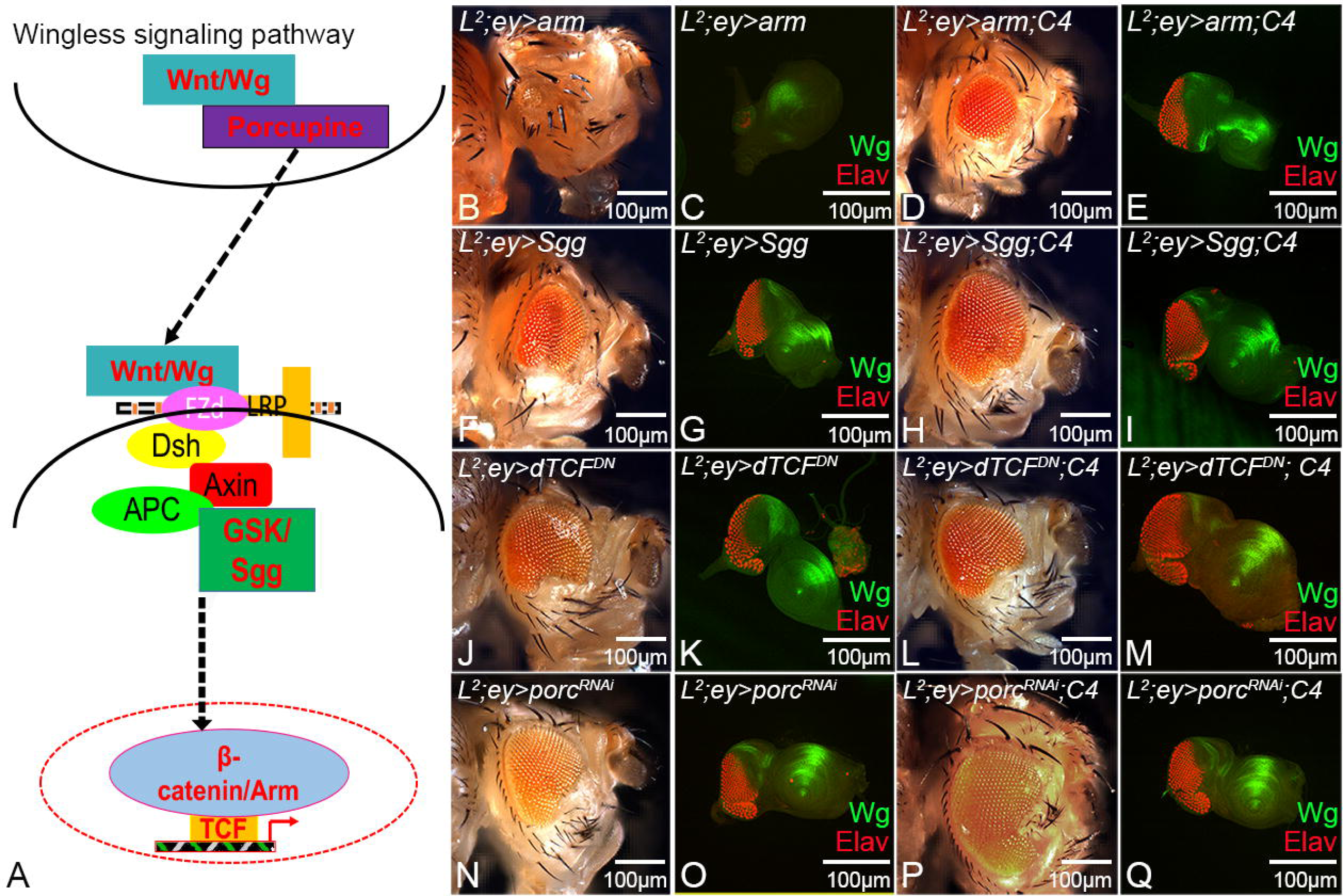
Modulating positive and negative regulators of Wg affect *L* mutant phenotype. **(A-Q)** Schematic presentation of Wg signaling pathway showing various members of the canonical pathway. **(B-E)** Activating Wg signaling in *L^2^* mutant eye background by misexpression of **(B, C)** *arm* alone (*L^2^*/+*; ey>arm*), **(D, E)** *arm* and *C4* (*L^2^*/+*; ey>arm; C4*). Note that *C4* misexpression with *arm (L^2^*/+*; ey>arm; C4)* in the eye rescues the *L^2^*/+*; ey>arm* reduced eye phenotype. **(F-M)** Blocking Wg signaling in *L^2^* mutant eye background by misexpression of **(F, G)** *sgg* alone (*L^2^*/+*; ey>sgg*), **(J, K)** *dTCF^DN^* alone (*L^2^*/+*; ey> dTCF^DN^*) results in rescue of *L^2^* loss-of-ventral eye phenotype. However, in *L^2^* mutant background misexpression of *C4* along with **(H, I)** *sgg* (*L^2^*/+*; ey>sgg; C4*) and **(L, M)** *dTCF^DN^* (*L^2^*/+*; ey> dTCF^DN^; C4*) significantly rescue the *L^2^*/+*; ey>sgg* and *L^2^*/+*; ey> dTCF^DN^* partial loss-of-ventral eye phenotype.**(N-Q)** Blocking transport of Wg morphogen in *L^2^* mutant eye background by misexpression of (N, O) *porc^RNAi^* alone (*L^2^*/+*; ey> porc^RNAi^*), (D, E) *porc^RNAi^* and *C4* (*L^2^*/+*; ey> porc^RNAi^; C4*). Note that *L^2^*/+*; ey> porc^RNAi^* exhibits weak rescue of loss-of-ventral eye phenotype whereas *C4* misexpression with *sgg (L^2^*/+*; ey> porc^RNAi^; C4)* in the eye rescues the *L^2^*/+*; ey> porc^RNAi^* small eye phenotype. All the images are displayed in same polarity as dorsal domain-towards the top, and ventral domain-towards the bottom. Scale bar= 100 μm.

Blocking Wg signaling by misexpressing *shaggy (sgg*), an antagonists of Wg signaling, in the eye (*ey>sgg,* Figure 6–figure Supplement 1E, F) (Hazelett et al., 1998; Singh et al., 2002; Singh et al., 2006) or along with *C4* in the wild-type background (*+*/+*; ey>sgg; C4,* Figure 6–figure Supplement 1G, H) does not affect the eye size. Misexpression of *sgg* in the *L^2^*/+ mutants (*L^2^*/+*; ey>sgg*), significantly rescues the loss-of-ventral eye phenotype in 27.3% of flies (Figure 6F, G, Figure 6–figure Supplement 2B and Table S1) (Singh et al., 2006). Similarly, misexpressing *C4* and *sgg* in the *L^2^*/+ mutant eye (*L^2^*/+*; ey>sgg; C4*) rescues the loss-of-ventral eye phenotype (Figure 6H, I) and rescue frequency increased dramatically from 27.3% to 71.5% (Figure 6–figure Supplement 2B and Table S1). The transcription factor dTCF is the downstream target of Wg signaling, and is downregulated by misexpression of the N-terminal deleted dominant-negative form of TCF (dTCF^DN^) (van de Wetering et al., 1997). Misexpression of dTCF^DN^ alone in the eye (*ey>dTCF^DN^*, Figure 6–figure Supplement 1I, J) or dTCF^DN^ with *C4* (*+*/+*; ey> dTCF^DN^; C4*, Figure 6–figure Supplement 1K, L) does not affect the eye size. Misexpressing *dTCF^DN^* in the eye of *L^2^*/+ mutants (*L^2^*/+*; ey>dTCF^DN^*) significantly rescues the loss-of-ventral eye phenotype in 21.8% of flies (Figure 6J, K, Figure 6–figure Supplement 2B and Table S1) (Singh et al., 2006). Similarly, misexpressing C4 and *dTCF^DN^* in the eye of *L^2^* mutants (*L^2^*/+*; ey> dTCF^DN^; C4*) rescues the loss-of-ventral eye phenotype (Figure 6L, M). As expected rescue frequency again dramatically increased from 21.8% to 70.6% (Figure 6–figure Supplement 2B and Table S1).

Our results demonstrate that misexpression of newt genes when Wg signaling is downregulated (using antagonists of Wg) in *L^2^* mutant background (*L^2^*/+*; ey> sgg; C4,* 71.5%) or (*L^2^*/+*; ey> dTCF^DN^; C4*, 70.6%) significantly increased the rescue frequency of the *L^2^* mutant phenotype. The rescue frequency obtained is greater than either misexpressing candidate gene *(L^2^*/+*; ey>C4,* 58.2%) alone or only misexpressing negative regulators (*L^2^*/+*; ey> sgg*, 27.3%) or (*L^2^*/+*; ey> dTCF^DN^*, 21.8%). The converse phenotypes were seen when Wg signaling was upregulated. This strongly suggest that newt gene(s) interact with the Wg signaling pathway to promote rescue of *L^2^* mutant phenotype.

Porcupine (Porc) is involved in post-translational modification of Wg nascent protein in the producing cell and thus facilitates its transport outside to the receiving cell. Wg ligand can bind to its receptor on the receiving cell (Manoukian et al., 1995; Swarup and Verheyen, 2012) thus modulating downstream pathway components (Figure 6A). We blocked Wg transport from the producing cell using *porc porc^RNAi^* and sampled its effect on *L^2^* mutant phenotype, and *L^2^* mutant phenotype where C4 is misexpressed. Misexpression of *porc^RNAi^* alone (*ey> porc^RNAi^*) (Figure 6–figure Supplement 1M, N), and along with C4 in the wild-type background (*+*/+*; ey> porc^RNAi^; C4*) (Figure 6–figure Supplement 1O, P) does not affect the eye size. Misexpressing *porc^RNAi^* in the *L^2^* mutant eye (*L^2^*/+*; ey> porc^RNAi^*) significantly rescues the loss-of-ventral eye phenotype in 20.8% of flies (Figure 6N, O, Figure 6–figure Supplement 2B and Table S1) whereas when *C4* is also misexpressed in the eye of *L^2^* mutants along with *porc^RNAi^* (*L^2^*/+*; ey> porc^RNAi^; C4*) rescued the loss-of-ventral eye phenotype (Figure 6P, Q). The rescue frequency increased dramatically from 20.8% to 68.1% (Figure 6–figure Supplement 2B, and Table S1). Thus, newt genes can modulate levels of Wg signaling to promote regeneration response in *L^2^* eye mutants.

## Discussion

*Notophthalmus viridescens* have enormous genome size (∼ c × 10^10^ bases), a long reproductive cycle, and have limited genetic tools that makes it difficult to use this model to ascertain the molecular mechanism(s) behind the function of its novel genes (Mehta and Singh, 2019a). One of the strategies can be to introduce these genes into models with array of genetic tools and where regeneration potential is lower as seen in models like *Drosophila, Bombyx mori* etc. (Gopinathan et al., 1998; Kango-Singh et al., 2001; Mehta and Singh, 2019a; Singh et al., 2007). The rationale is to test the efficacy of these novel newt genes in triggering regeneration response in these model systems with lower regeneration potential. We and others have introduced human, plant and other vertebrate genes by transgenic approaches in *Drosophila* (Deshpande et al., 2019; Gogia et al., 2020b; Hughes et al., 2012; Sarkar et al., 2018; Tare et al., 2011). These transgenic flies have been utilized for targeted expression of foreign genes in flies using a targeted misexpression approach (Brand and Perrimon, 1993). The successful misexpression of the unique newt genes (vertebrates) in *Drosophila* using the same targeted misexpression approach has been reported earlier (Mehta et al., 2019; Mehta and Singh, 2019b).

To test the regeneration potential of these novel newt genes, we took a distinctive approach of using *Drosophila* model because the signaling pathways, which are involved in regeneration and/or tissue growth, patterning and development are evolutionarily conserved across the species (Kango-Singh and Singh, 2009; Mehta and Singh, 2019a; Wang and Hu, 2020). Even though humans *(Homo sapiens),* which evolved approx. 200,000 years ago (Stringer, 2016) and are 541 million years apart from *Drosophila* still share 75% of genes with flies (Pandey and Nichols, 2011). Despite the gap of 250 million years between newts and *Drosophila*, our studies suggest that the proteins encoded by unique newt genes are functional in *Drosophila* (Figure 1). Therefore, it is plausible that the pathways that newt genes can modulate in *Drosophila* might share parallels with their mechanism of action in newts. We found that the five newly identified genes from newt, can (i) rescue *L* mutant eye phenotype (early eye mutant) as well as GMR-*hid* GMR-GAL4 (late retinal degeneration) phenotype, (ii) promote cell proliferation, and downregulate cell death to restore loss-of-ventral eye phenotype, and (iii) downregulate Wg signaling (Figure 7). Our studies clearly demonstrate the regeneration potential of these novel newt genes in triggering rescue response in *Drosophila* model along the temporal axis.

**Figure 7.**
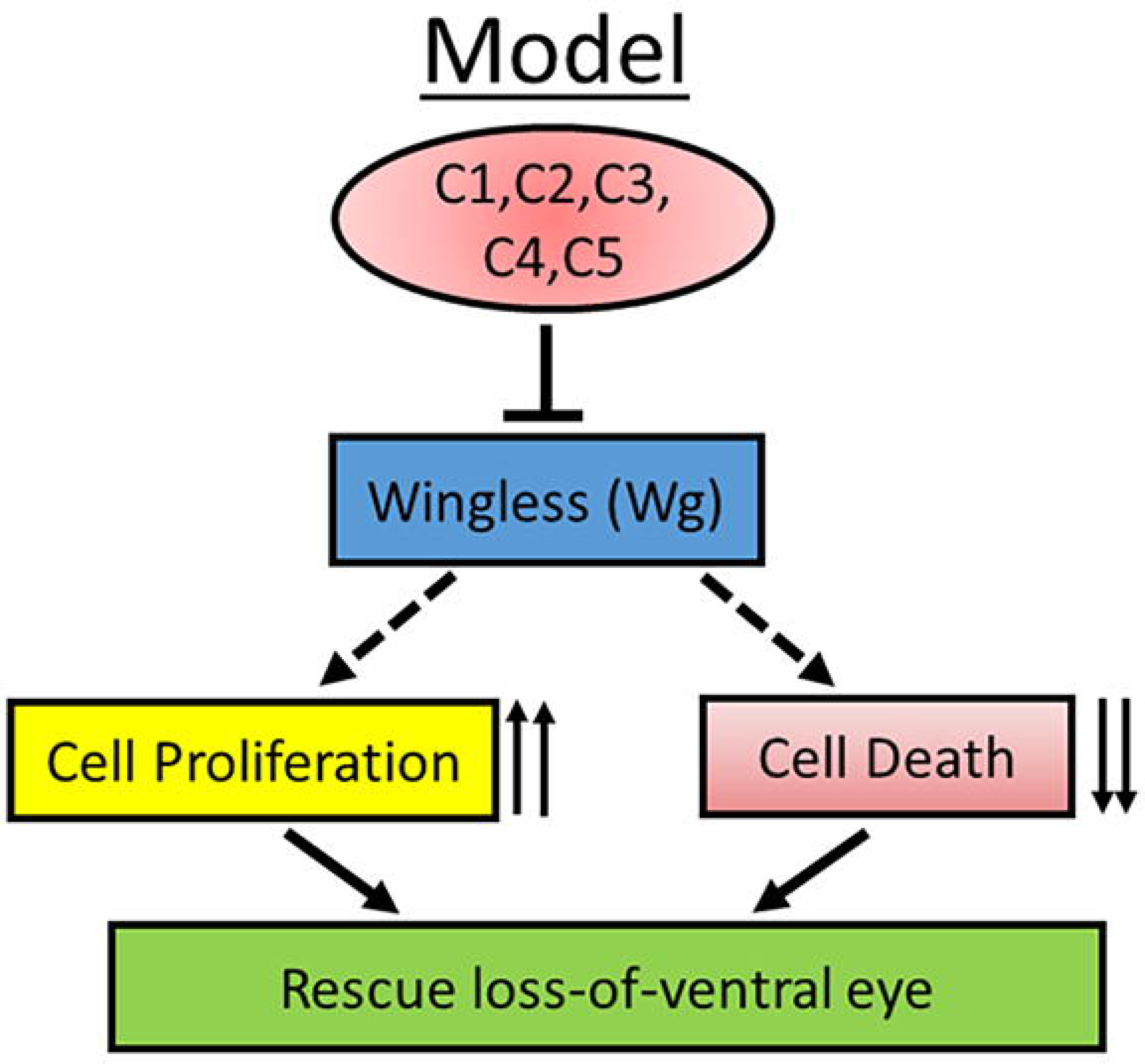
Model of newt genes function during development. These novel newt genes (C1-C5) can negatively regulate evolutionarily conserved Wg /Wnt signaling pathway in *Drosophila*, promote cell proliferation, and block cell death, which together results in the rescue of *L^2^* mutant loss-of-ventral eye phenotype. This regenerative ability of these novel newt genes extend from early eye development to late larval eye and adult eye.

### Newt genes promote cell proliferation to restore *Drosophila* eye mutant phenotype

Regeneration involves the formation of a blastema, a proliferating zone of undifferentiated cells to restore the missing structures (Brockes and Kintner, 1986; Mehta and Singh, 2019a; Sanchez Alvarado and Tsonis, 2006). Furthermore, a network of newt gene(s) trigger proliferation response to promote lens regeneration (Eguchi and Shingai, 1971; Grogg et al., 2005; Looso et al., 2013; Madhavan et al., 2006) A similar proliferating response was observed when newt genes were misexpressed in the *Drosophila L^2^* mutant eye (Figure 3C, and Figure 3–figure Supplement 1). *L^2^* mutant flies exhibit a loss-of-ventral eye phenotype, and misexpressing newt genes causes 4.2-5.8 fold increase in number proliferating cells (Figure 3, 7 and Figure 3–figure Supplement 1), which clearly mimics the classic mechanism of epimorphic regeneration in the newt lens.

In our study, we found that newt genes can also modulate cell death in *Drosophila* developing eye. When newt genes were misexpressed in the *L^2^* mutant background (*L^2^*; *ey*>*newt gene*), the rate of cell death was lower as compared to the *L^2^* mutant (*L^2^*; *ey*-GAL4) eye (Figure 3E, F, H and Figure 3–figure Supplement 1H-L, N), but minimally higher compared to the positive control (*ey*-GAL4) (Figure 3F, G, H and Figure 3–figure Supplement 1I-N). A minimal rate of apoptosis is observed during the newt lens regeneration (Tsonis et al., 2004). Our data suggest that newt genes hijack the evolutionarily conserved signaling pathway(s) in *Drosophila* to regulate cell proliferation, and cell death. It is a strong possibility that the same pathways may promote regeneration in newts.

### Newt genes tightly regulate Wg signaling pathway in *Drosophila*

We screened for the various signaling pathways using the candidate approach as well as compared them to our transcriptomic analysis (Mehta et al., 2019). Among the various genetic modifiers of the *L^2^* mutant phenotype, we found *wg* as one of the possible candidate. Wg, a member of the evolutionarily conserved Wg/Wnt pathway, has been previously reported to promote regeneration, growth, development etc. (Harris et al., 2016; Schubiger et al., 2010; Singh et al., 2012b; Singh et al., 2018; Smith-Bolton et al., 2009; Yokoyama et al., 2007). Misexpresion of newt genes in *Drosophila* wild-type background modulate both agonists of Wg/Wnt signaling like *frizzled (fz),* and *microtubule star,*(*mts*) and antagonists like *Casein Kinase II*β (*CKII*β), and *sinaH* (Figure 4–figure Supplement 1). This modulation in the levels of both antagonists as well as agonists can be a homeostatic reaction shown by the *Drosophila* to normalize expression levels of Wg/Wnt signaling pathway. As a result, changes in the eye growth after misexpressing newt genes in the wild-type was non-significant when compared to the wild-type control (*ey*-GAL4) (Figure 1). We did not observe any lethality, developmental defects etc.

Interestingly, when newt genes were misexpressed in *L^2^* mutant background, there was significant downregulation of Wg/Wnt expression (Figures 4, 7, and Figure 4–figure Supplement 2), which resulted in rescue of the loss-of-ventral eye phenotype. It is known that overexpressing Wnt, and its receptor Fz promote lens regeneration in newts, whereas blocking Wnt pathway by expressing Dkk1 inhibits lens induction (Hayashi et al., 2006). Therefore, our study may have parallels with the regeneration mechanism in newts.

Furthermore, we found that when Wg pathway components are modulated in *Drosophila* along with the newt genes, it not only rescue the *L^2^* mutant loss-of-ventral eye phenotype (Figure 6) but can also rescue the eye-loss caused by the activation of the Wg signaling pathway in the wild-type background (Figure 6–figure Supplement 1). Hence, our data strongly suggest that the newt genes can modulate Wg signaling pathway in loss-of-ventral eye background. Interestingly, these newt genes did not show any phenotype when they were misexpressed alone suggesting that regeneration response is triggered only to the cues generated due to tissue damage.

### Newt proteins exhibit non-cell-autonomous response

Genetic mosaic experimental approach, which resulted in the generation of clone(s) of retinal neurons expressing high levels of newt genes within the *L^2^* mutant eye disc. These experiments showed that clones expressing high levels of newt genes can rescue the *L^2^* mutant phenotype, however the rescue was not limited to the clone itself and spread into the neighboring cells where newt genes were not misexpressed. This non-cell-autonomous rescue of the missing tissue is possible only if the newt proteins are getting secreted from one cell to another and/or newt proteins are acting on the signaling molecules. These newt proteins have signal peptides (Looso et al., 2013), which can facilitate its transport from one cell to another. Similarly, we have reported that these genes interact with Wg (Figure 4), which is a morphogen that can have a long distance effect in the *Drosophila* eye (Zecca et al., 1996a).

It will be interesting to investigate, (a) if newt genes can regulate Wg signaling pathway to promote regeneration in other tissues of *Drosophila* e.g. wing, leg etc., and under the background of other congenital mutations (Schubiger et al., 2010; Smith-Bolton et al., 2009), (b) if these proteins can rescue/regenerate missing tissues/organs in other animals with low regenerative potentials. The pathways that have roles in regeneration, growth, development are evolutionarily conserved throughout the animal kingdom (Mehta and Singh, 2019a). For example, the Wg/Wnt signaling pathway, is one such evolutionarily conserved pathway that apart from promoting regeneration in hydra, planaria, newts, zebrafish, etc. has also been found to promote cell proliferation during regeneration of mammalian muscles, liver, and bone (Polesskaya et al., 2003; Sodhi et al., 2005; Zhong et al., 2006). Therefore, it raises the possibility that genetically engineering genes from the highly regenerative animals that show regeneration in a precisely controlled fashion might systematically regulate evolutionarily conserved pathways in a foreign host to promote tissue regeneration and/or cell survival.

## Materials and methods

### Animals

Handling and operations of *Notophthalmus viridescens* (newts) have been described previously (Mehta et al., 2019; Sousounis et al., 2013). For our studies, *Notophthalmus viridescens* (newts) were purchased from Charles Sullivan Inc. newt farm. All procedures involving animals were approved by the University of Dayton Institutional Animal Care and Use Committee (IACUC; Protocol ID: 011-02). Newts were anesthetized in 0.1% (w/v) ethyl-3-aminobenzoate methanesulfonic acid (MS222; Sigma) dissolved in phosphate buffered saline. The surgical procedures were performed in anesthetized with MS222 newts, and appropriate procedures were undertaken to alleviate pain and distress while working with newts.

### Generation of transgenic flies

The design and generation of transgenic flies (UAS-C1, UAS-C2, UAS-C3, UAS-C4, and UAS-C5) used in this study have been described (Mehta et al., 2019). To validate the successful insertion of newt genes in *Drosophila* by targeted misexpression approach, we generated transgenic fly like UAS-C1-5-V5 with 14 aa long V5-epitope sequence tagged towards the 3’ end of the C4 ORF. The complete gene fragment (5’-C4-V5-3’) was synthesized, and cloned into the XbaI and XhoI restriction sites in pUAST-attB plasmid. The pUAST-attB plasmid containing both the transgene and donor sequence (attB) was co-injected along with ⎞C31 integrase mRNA into attP-containing recipient embryos, resulting in site-specific insertion of the transgene at Chromosome III (Fish et al., 2007; Groth et al., 2004). The integration and expression of transgene was verified by the western blots using V5 antibody.

### *Drosophila* stocks

In this study, we used transgenic flies UAS-C1, UAS-C2, UAS-C3, UAS-C4, UAS-C5, and UAS-C1-5-V5). Other fly stocks used in this study are described in Flybase (http://flybase.bio.indiana.edu). We used *Tub-*GAL4*/S-T* (Yang et al., 2019)*, L^2^/CyO; ey-*GAL4*/ey-*GAL4, *L^2^/C4; ey-*GAL4*/ey-*GAL4*, UAS-GFP* (Singh et al., 2005; Singh and Choi, 2003; Singh et al., 2006). *DE-*GAL4*/CyO; UAS-GFP/Tb* (Morrison and Halder, 2010), *UAS-wg*-*GFP* (Azpiazu and Morata, 1998), *UAS-arm (Zecca et al., 1996b), UAS-sgg* (Hazelett et al., 1998)*, UAS-dTCF^DN^* (van de Wetering et al., 1997) *, UAS-p35* (Hay et al., 1994).

### Targeted misexpression studies

We used the GAL4/UAS system for targeted misexpression studies (Brand and Perrimon, 1993). All GAL4/UAS crosses were performed at 25°C. In order to misexpress the UAS-transgenes, we used *tub* –GAL4 (Yang et al., 2019) to ubiquitously misexpress genes*, ey-*GAL4 (Hazelett et al., 1998) to misexpress transgene in the developing eye, and *DE-*GAL4 (Morrison and Halder, 2010) that drives transgene in the dorsal domain of the developing eye.

### Genetic mosaic analysis

Clones were generated using the FLP/FRT system of mitotic recombination (Xu and Rubin, 1993). To generate clones of C4 *in L^2^*-mutant eye, *yw* hsFLP; Act>y+>GAL4, UAS-*GFP*/*CyO* females were crossed to *L^2^/CyO;* UAS*-C4/Tb* males. C4 is expressed in all the GFP+ cells of eye-antennal imaginal disc.

### Immunohistochemistry

Eye-antenna discs were dissected from wandering third instar larvae and stained following the standard protocol (Singh et al., 2006). The eye imaginal discs were dissected in cold 1X solution of Phosphate Buffered Saline (PBS) and fixed using 4% p-formaldehyde (Electron Microscopy Sciences, Catalog no. 15710). Fixed tissues were washed using PBS-T [0.2% Triton X-100 (Sigma, Catalog no. T8787) in 1XPBS] and immunostained using primary antibodies: mouse anti-Phospho histone-3 (1:200) (Sigma-Aldrich); Rabbit anti-*Drosophila* caspase-1 (DCP-1) (1:100) (Sigma-Aldrich); rat anti-Elav (1:100) (Developmental Studies Hybridoma Bank); mouse anti-Wg (1:100) (Developmental Studies Hybridoma Bank), and goat anti-Hth (a gift from H. Sun and R. Mann). Secondary antibodies (Jackson Laboratories) used were goat anti-rat IgG conjugated with Cy5 (1:200); donkey anti-rat IgG conjugated to FITC (1:200); donkey anti-rat IgG conjugated to Cy5 (1:200), donkey anti-mouse IgG conjugated to Cy3 (1:200); donkey anti-goat IgG conjugated to Cy3 (1:200), and goat anti-rabbit IgG conjugated with Cy5 (1:200). The discs were mounted in Vectashield (Vector labs, H1000) and imaged using Olympus Fluoview 1000 microscope.

### Western Blot

In this study, two different western blots were run. For blot 1, protein samples were prepared from (*n* = ∼10) the wandering third instar larvae of transgenic fly UAS-C4-V5 as experimental sample, and *ey*-GAL4 was used as control. For blot 2, protein samples were prepared from (*n* = ∼25) the adult heads of *ey*-GAL4, *L^2^; ey*-GAL4 and *L^2^; ey*>*C4*. Protein samples were isolated by standardized protocol (Gogia et al., 2017), ran on 10% gel, and then transferred onto nitrocellulose membranes. For blot 1, the membrane was then incubated overnight at 4°C with primary antibodies: 1∶15000 rabbit anti-V5 antibody (SIGMA, Cat. # V8137), and 1:12000 mouse anti-α-Tubulin antibody (SIGMA, Cat. # T5168). Followed by 1h incubation with secondary antibodies: horseradish peroxidase conjugated goat anti–rabbit IgG (1:5000 dilution) secondary antibody, and goat anti-mouse IgG-HRP (1:5000 dilution) (Santa Cruz Biotechnology, Cat. Sc-2005). For blot 2, the membrane was then incubated overnight at 4°C with primary antibody 1:12000 mouse anti-α-Tubulin antibody (SIGMA, Cat. # T5168). Followed by 1h incubation with secondary antibody: horseradish peroxidase conjugated goat anti-mouse IgG-HRP (1:5000 dilution) (Santa Cruz Biotechnology, Cat. Sc-2005). The blot was then stripped and then incubated overnight with primary antibody 1∶1000 mouse anti-Wingless antibody (DSHB, Cat. # 4D4). Followed by 1h incubation with secondary antibody 1:5000 horseradish peroxidase conjugated goat anti-mouse IgG-HRP (Santa Cruz Biotechnology, Cat. Sc-2005). The signal in both the blots were detected using Super Signal West Dura Extended Duration Substrate (ThermoFisher Scientific, Cat. #34076). Images were captured using the UVP Imaging System. Wg levels were quantified and normalized by using Licor Image Studio lite 5.2 software and this was followed by statistical analysis.

### TUNEL assay for detection of cell death

Apoptosis was detected by using Terminal deoxynucleotidyl transferase-mediated dUTP nick-end labeling (TUNEL) assay (Tare et al., 2011; White et al., 2001). TUNEL assay is based on the detection of fragmented DNA ends, which are characteristic of apoptotic cells. A deoxynucleotidyl transferase (TdT) is used to add fluorescently labeled nucleotides to free 3’-OH DNA ends in a template-independent manner. The modified nucleotides can then be detected by fluorescence microscopy. Eye-antennal discs, after secondary-antibody staining were blocked in 10% normal goat serum in phosphate buffered saline with 0.2% Triton X100 (PBT) for an hour. After blocking, samples were incubated in 0.1 M sodium citrate (Fisher Scientific) and 10% TritonX-100 for 30 minutes in dark at 65°C. Samples were washed and incubated in TUNEL dilution buffer (Roche Diagnostics, Catalog no. 11966006001) for 10 minutes at room temperature. Followed by labeling with 50% TUNEL labeling solution for 30 minutes (Roche Diagnostics, Catalog no. 12156792910) in dark at room temperature, and finally samples were subjected to TdT enzyme solution (Roche Diagnostics) for 2 hours at 37°C. After which samples were washed with PBS-T and mounted in Vectashield (Vector labs, Catalog no. H1000).

### Bright Field imaging

Adult flies were selected and were transferred to −20°C for two hours, followed by removal of legs and wings. The legs and wings were removed in order to get a clear view of the compound eye. The Bright field pictures of adult heads were taken using a Zeiss Apotome Imager Z1 (Wittkorn et al., 2015). Eye size was measured using ImageJ software tools (http://rsb.info.nih.gov/ij/). To normalize the eye size values and neglect any variations caused by overall different head size we took ratio of eye size to the head size for each individual picture. Then relative ratio was calculated for all the treatments, and the value reported is the relative fold change for each experimental treatment compared to the control.

### Statistical analysis

Statistical analysis was performed using student’s t-test for independent samples. Sample size used for the calculation was five in number (n=5), otherwise specified. Statistical significance was determined with 95% confidence (p<0.05).

## Acknowledgments

We thank Bloomington *Drosophila* Stock Center (BDSC) for *Drosophila* strains, and the Developmental Studies Hybridoma Bank (DSHB) for antibodies. We would like to thank to Katia Del Rio-Tsonis, Labib Rouhana, Madhuri Kango-Singh, Meghana Tare, Neha Gogia, Shilpi Verghese, Pothitos Pitychoutis, Deepika K Sodhi for critical comments on the manuscript. We also thank Justin Kumar, Y. Henry Sun, and Kyung Ok Cho for gift of fly strains and antibodies. Confocal microscopy was supported by core facility at University of Dayton.

## Funding resources

A.S. is supported by NIH1R15GM124654-01 from NIH, Schuellein Chair Endowment Fund and STEM Catalyst Grant from the University of Dayton.

---

**Figure 1–figure Supplement 1. Validation of newt genes misexpression in the *Drosophila*.** To validate the misexpression of newt gene in *Drosophila* we tagged candidate 4 with **(A)** V5 epitope **(B)** Generated transgenic fly containing *C4-V5* fused gene **(C)** misexpressed in the F1 progeny and **(D)** performed western blotting against V5 epitope. **(A)** The 42 bp long V5 tag is fused towards the 3’ end of the 501 bp long open reading frame (ORF) of newt gene (*C4*). **(B)** The ORF comprising both *C4* and *V5* fused together as 501+42= 543 bp long (‘C4-V5’) is cloned into the pUAST-attB plasmid. The plasmid is microinjected at a specific site (attP) on the III chromosome in the *Drosophila* to generate a transgenic fly strain: UAS–*C4-V5*. **(C)** *Drosophila* strain trapped with *tub*-GAL4 enhancer is mated with transgenic *Drosophila* (*UAS–C4-V5*) to drive *C4-V5* misexpression ubiquitously in the (F1) progeny (*tub>C4-V5*). The protein is extracted from the third instar larvae of F1 progeny (*tub>C4-V5*), n=10. **(D)** C4-V5 fused protein is 166 aa (C4) + 14 aa (V5)= 180 aa long, which is equivalent to molecular weight of 19.8 kilodalton (kD). The Blot shows the expression of fused candidate protein (C4-V5) in *Drosophila melanogaster.* The samples were loaded in the following sequence: Lane 1- *tub*>C4-V5, Lane 2- *ey*>GAL4 (Control), Lane 3- Molecular weight marker. Alpha tubulin is used as a loading control. Molecular weight of alpha tubulin is 55 kD. The samples were treated with anti-V5 tag antibody, and anti-α tubulin antibody. The presence of V5 band (of molecular weight 19.8 kD) in *tub*>*C4-V5* lane suggests that *C4* (newt gene) is expressed in *Drosophila melanogaster* using targeted misexpression approach.

**Figure 1–figure Supplement 2. Rescue frequency of loss-of-ventral eye phenotype upon misexpression of respective *newt genes* in *L^2^mutant* background.** Rescue frequency was about 40.83% in *C1*, 37.7% in *C2*, 49% in *C3*, 58.2% in *C4*, and 38.7% in *C5* respectively. All bar graphs show rescue frequency as average between 3 repetitions. Error bars show standard deviation, and numbers above the error bar show rescue frequency in percentage. Two hundred flies were counted per repetition (200 × 3= 600) for calculating the frequency for each sample. *L^2^*-mutant shows 0% penetrance.

**Figure 1–figure Supplement 3. Unique newt candidate genes can also promote rescue in later stages of eye development. (A-C)** Bright field adult eye pictures **(A)** Wild-type adult eye as positive control (*ey-*GAL4) **(B)** GMR-*hid*,GMR-GAL4 (GMR>*hid*) adult eye as negative control. Misexpression of *hid* results in a “No-eye” or highly reduced eye phenotype. **(C)** Misexpression of newt gene (*C4*) in GMR-*hid*, GMR-GAL4 (GMR>*hid;C4)* background result in a significant rescue of the “No-eye” phenotype. Thus, confirming that newt genes can also promote rescue in later stages of eye development. All the images are displayed in same polarity as dorsal domain-towards top, and ventral domain-towards bottom. Scale bar= 100 μm.

**Figure 1–figure Supplement 4. All the members of the unique newt gene family induce cell proliferation, and block cell death to promote rescue of *L^2^* mutant loss-of-ventral eye phenotype.** Testing other members of the unique newt gene family we found same trend in upregulation of (A-G) cell proliferation, and (H-N) downregulation in cell death. (A-F & A′-F′) eye discs showing PH-3 as a marker to calculate mitotic index (A) *L^2^* mutant eye disc (B-E) *L^2^* mutant eye disc under the background of which newt genes (B*) C1,* (C) *C2*, (D) *C3*, and (E) *C5* are misexpressed respectively (F) Wild-type (*ey*-GAL4) eye disc as positive control. (G) bar graph showing relative increase in the mitotic index (MI) when newt genes are misexpressed in *L^2^* mutant background. These results show that all the *newt genes* behave similarly and induce cell proliferation to restore the missing cells. And thereby significantly rescuing the *L^2^* mutant’s loss-of-ventral eye phenotype. (H-M & I′-M′) Eye discs showing TUNEL staining as a marker for cell death assay in (H) *L^2^* mutant eye disc (J-N) *L^2^* mutant eye disc where newt genes (I) *C1*, (J) *C2*, (K) *C3,* and (L) *C5* are misexpressed respectively (M) Wild type (*ey*-GAL4) eye disc as positive control. (N) Bar graph showing relative decrease in the cell death when newt genes are misexpressed in the *L^2^*-mutant background. These results together show that the *newt genes* apart from upregulating cell division also downregulate number of dying cells in order to restore, and rescue *L^2^* mutant loss-of-ventral eye phenotype. Statistical analysis was performed using student’s t-test for independent samples. Sample size used for the calculation was five in number (n=5). Statistical significance was determined with 95% confidence (p<0.05). All bar graphs show relative fold change in cell proliferation and cell death as average between 5 samples. Error bars show standard deviation, and symbols above the error bar ** signify p value <0.005, *** signify p value <0.0005 and ns signify p value > 0.05 respectively. All the images are displayed in same polarity as dorsal domain-towards top, and ventral domain-towards bottom. Scale bar= 100 μm.

**Figure 1–figure Supplement 5. Downregulation in dying cells by misexpression of newt genes in *L^2^*-mutant background. (A-G)** Eye discs marked by cleaved *Drosophila* DCP-1 antibody. *Drosophila* cell death caspase (DCP-1) is *Drosophila* homologue of human caspase 3 that marks dying cells. **(A**′**-G**′) are single channel images showing number of dying cells in respective eye discs. Results again show that misexpressing newt genes downregulate number of dying cells in *L^2^* mutant background. All the images are displayed in same polarity as dorsal domain-towards top, and ventral domain-towards bottom. Scale bar= 100 μm.

**Figure 1–figure Supplement 6. Comparison of rescue of *L^2^* mutants by baculo-virus P35 mediated downregulation of cell death with misexpression of newt gene *C4*. (A)** *L^2^* mutant eye (*L^2^; ey-*GAL4) **(B)** *L^2^* mutant adult eye that contains *p35* as a transgene (*L^2^*/+*;p35*) **(C)** *L^2^* mutant adult eye in which p35 is misexpressed using ey-GAL4 (*L^2^*/+*; ey>p35*). **(D)** *L^2^* mutant adult eye in which *C4* newt gene is misexpressed (*L^2^*/+*; ey>C4*). **(E)** Wild type adult eye (*ey*-GAL4) as positive control. **(F)** Phenotype strength between the samples. **(G)** Comparison of loss-of-ventral eye phenotype rescue frequency induced by *p35* misexpression (*L^2^*/+*; ey>p35*) with the one induced by *C4* (newt gene) misexpression (*L^2^*/+*; ey>C4*). Both misexpressed in the *L^2^* mutant background. Note that *L^2^* mutants loss-of-ventral eye phenotype rescued by anti-apoptotic gene, *p35* (*L^2^*/+*; ey>p35*) are weaker in strength, and rescue frequency is significantly less when compared to the *L^2^* mutant fly eye in the background where newt gene is misexpressed (*L^2^*/+*; ey>C4*). Statistical analysis was performed using student’s t-test for independent samples. Statistical significance was determined with 95% confidence (p<0.05). In case of **(F)** phenotype strength all bar graphs show relative fold change in the eye size as average between 5 samples. In case of **(G)** rescue frequency percentage all bar graphs show rescue frequency as average between 3 repetitions. Two hundred flies were counted per repetition for calculating the frequency for each sample. Error bars show standard deviation and symbols above the error bar * signify p value <0.05, and ** signify p value <0.005 respectively. *L^2^*-mutant shows 0% penetrance. All the images are displayed in same polarity as dorsal domain-towards top, and ventral domain-towards bottom. Scale bar= 100 μm.

**Figure 1–figure Supplement 7. Misexpression of newt genes affect evolutionarily conserved Wg pathway components.** Next generation RNA sequencing for wild-type *Drosophila* samples collected at third-instar larval stage under the background of which novel *newt genes* were misexpressed revealed that **(A)** Wingless/ Wnt was one of the important evolutionary conserved pathway that was reported to be differentially regulated by misexpressing newt genes ubiquitously in the developing *Drosophila*. **(B)** The components of the Wingless/ Wnt pathway that are significantly affected include (1) *fz* (principal receptors for the Wnt family of ligands) showed 4-fold downregulation, (2) *microtubule star* (*mts*) (encodes the catalytic subunit of protein phosphatase 2A) showed 9-fold downregulation, (3) *casein kinase II beta* (*CKII*β) (component of β catenin degradation complex) showed 7 fold downregulation, and (4) *sina homologue* (*sinaH*) (codes for E3 ubiquitin-protein ligase) showed 12 fold downregulation.

**Figure 1–figure Supplement 8. All the newt gene family members’ downregulate Wg to promote rescue of *L^2^*-mutant loss-of-ventral eye phenotype. (A-F)** eye disc showing wg staining in green, and Elav in red **(A**′**-F**′) are single filter confocal images showing Wg staining as grey image. **(G, G**′) Bar graphs displaying intensity plots for Wg expression of representative samples. Results clearly show the same trend of significant downregulation in Wg expression when newt genes **(C, C**′) *C1*, **(D, D**′) *C2*, **(E, E**′) *C3*, **(F, F**′) *C5* are misexpressed under the *L^2^* mutant background compared to the **(B, B**′) *L^2^* mutant fly **(A, A**′) is a wild-type eye disc showing normal level of Wg staining. Statistical analysis was performed using student’s t-test for independent samples. Statistical significance was determined with 95% confidence (p<0.05). All bar graphs show integrative density as a scale to measure Wg intensity for each sample represented as the average between 5. Error bars show standard deviation, and symbols above the error bar * signify p value <0.05, and ** signify p value <0.005 respectively. All the images are displayed in same polarity as dorsal domain-towards top, and ventral domain-towards bottom. Scale bar= 100 μm.

**Figure 1–figure Supplement 9. Percentage rescue frequency of *L^2^* mutant loss-of-ventral eye phenotype.** Rescue frequency was about (avg) 58.2% when *C4* alone is misexpressed in the *L^2^* mutant background. Rescue frequency generated by *C4* is low when *wg* is misexpressed both in wild-type background (*+*/+*; ey>wg-GFP,* rescue frequency= 16.8), and under *L^2^*-mutant background (*L^2^*/+*; ey>wg-GFP; C4,* rescue frequency= 21). All bar graphs show rescue frequency as average between 3 repetitions. Error bars show standard deviation. Two hundred flies were counted per repetition for calculating the frequency for each sample. *L^2^* mutant shows 0% penetrance.

**Figure 1–figure Supplement 10. Modulation of positive and negative regulators of Wingless under wild type background.** Misexpression of **(A, B)** *arm* alone (*+*/+*; ey>arm*) **(C, D)** *arm* and *C4* (*+*/+*; ey>arm;C4*) **(E,F)** *sgg* (*+*/+*; ey>sgg*) **(G, H)** *sgg* and *C4* (+/+*; ey>sgg;C4*) **(I,J)** dTCF^DN^ (*+*/+*; ey>* dTCF^DN^) **(K, L)** dTCF^DN^ and *C4* (+/+*; ey>* dTCF^DN^;*C4*) **(M, N)** Porc^RNAi^ (*+*/+*; ey>* porc^RNAi^). **(O, P)** Porc^RNAi^ and *C4* (*+*/+*; ey>* porc^RNAi^; *C4*). Misexpression of positive regulator of Wg like *arm* cause eye-loss phenotype, and misexpressing *newt genes* e.g., *C4* interact with the positive regulator of Wg pathway e.g., *arm* causing partial rescue of eye-loss phenotype in the wild type background. Negative regulators of Wg pathway nor the misexpression of *newt gene* along with the negative regulators of Wg generates eye growth or any weird phenotype. Signifying that *newt genes* do not cause overgrowth, and they promote phenotypic transition effect only when there is the need to rescue eye-loss phenotype. Elav staining is shown in red. Wg in green. All the images are displayed in same polarity as dorsal domain-towards the top, and ventral domain-towards the bottom. Scale bar= 100 μm.

**Figure 1–figure Supplement 11. Percentage rescue frequency of *L^2^* mutant loss-of-ventral eye phenotype. (A)** Rescue frequency was about (avg) 58.2% when *C4* alone is misexpressed under the *L^2^*-mutant background. Rescue frequency generated by *C4* is greater compared to when *C4* is misexpressed along with *arm* both in the wild type background (*+*/+*; ey>arm,C4,* rescue frequency= 21), and in *L^2^* mutant background (*L^2^*/+*; ey>arm; C4,* rescue frequency= 22.8). **(B)** In case of negative regulators misexpressed under the *L^2^*-mutant background rescue frequency is (*L^2^*/+*; ey> sgg,* rescue frequency=27.3), (*L^2^*/+*; ey>* dTCF^DN^, rescue frequency=21.8), (*L^2^*/+*; ey>* porc^RNAi^, rescue frequency=20.8). All the values are below the rescue frequency obtained when *C4* alone is misexpressed (*L^2^*/+*; ey>* C4, rescue frequency=58.2). However, rescue frequency dramatically increases when *C4* is added to the negative regulators, (*L^2^*/+*; ey>sgg,* rescue frequency=71.5), (*L^2^*/+*; ey>* dTCF^DN^, rescue frequency=70.6), (*L^2^*/+*; ey>* porc^RNAi^, rescue frequency =68.1)). Signifying a strong interaction between newt gene and Wg signaling pathway components. All bar graphs show rescue frequency as average between 3 repetitions. Error bars show standard deviation. Two hundred flies were counted per repetition for calculating the frequency for each sample. *L^2^* mutant shows 100% penetrance.

**Supplementary Table 1:**
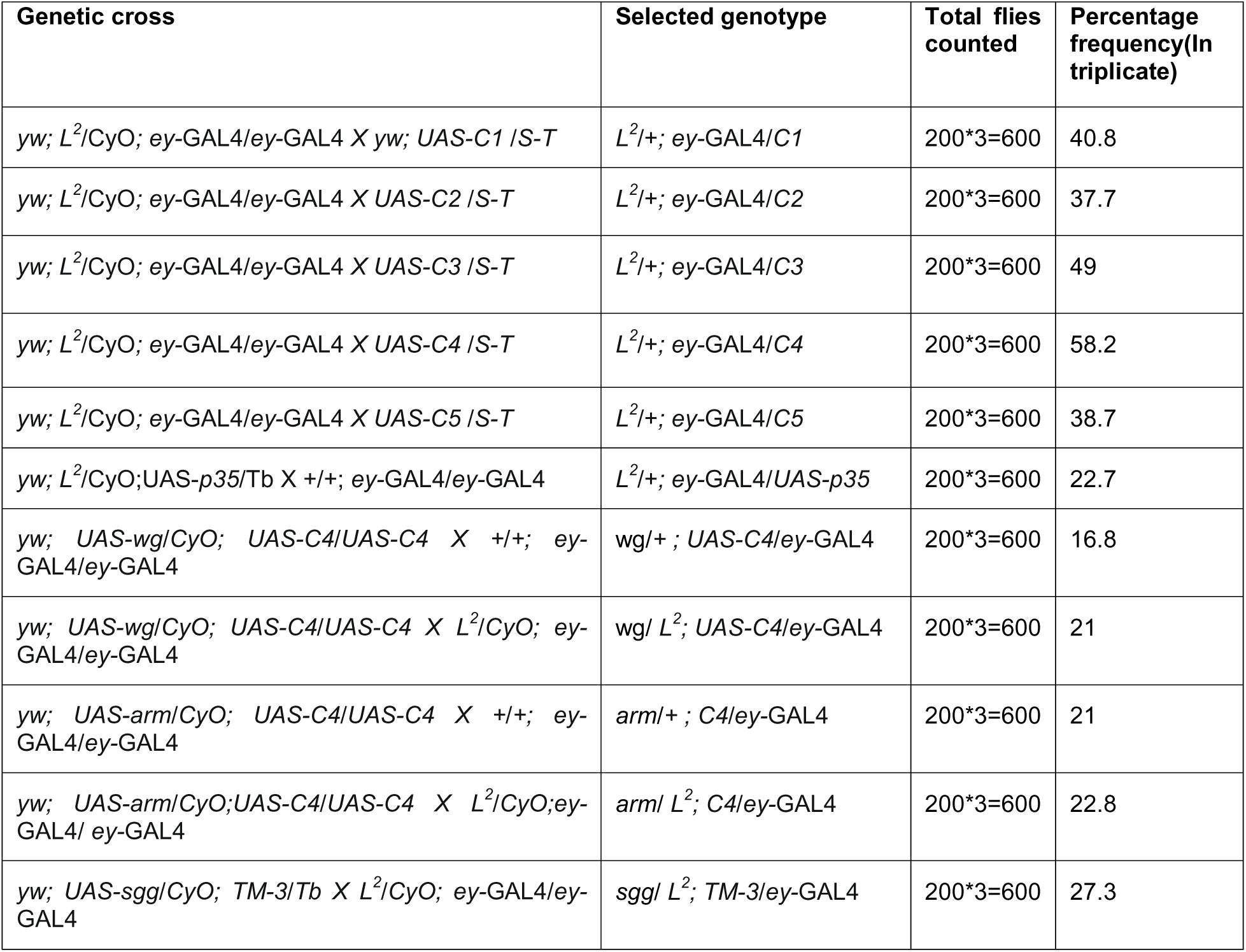
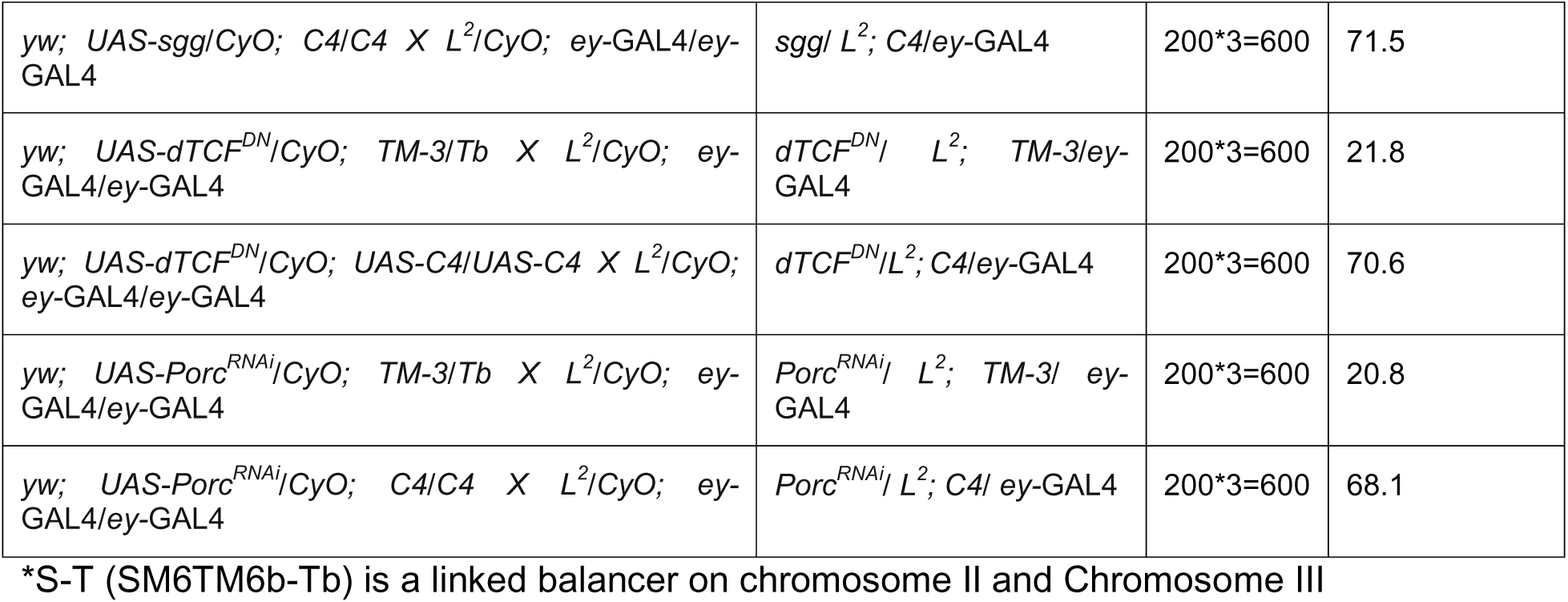
Rescue frequency calculated for respective Drosophila strains, and genetic crosses involved to achieve selective genotype.

