## Supplementary information for "Novel newt regeneration genes regulate Wingless signaling to restore patterning in *Drosophila* eye"

**Supplementary Table 1: Rescue frequency calculated for respective Drosophila strains, and genetic crosses involved to achieve selective genotype.**

| **Genetic cross** | **Selected genotype** | **Total flies counted** | **Percentage frequency(In triplicate)** |
| --- | --- | --- | --- |
| *yw; L^2^*/CyO*; ey-*GAL4/*ey-*GAL4 *X yw; UAS-C1 /S-T* | *L^2^*/+*; ey-*GAL4*/C1* | 200*3=600 | 40.8 |
| *yw; L^2^*/CyO*; ey-*GAL4*/ey-*GAL4 *X UAS-C2 /S-T* | *L^2^*/+*; ey-*GAL4*/C2* | 200*3=600 | 37.7 |
| *yw; L^2^*/CyO*; ey-*GAL4*/ey-*GAL4 *X UAS-C3 /S-T* | *L^2^*/+*; ey-*GAL4*/C3* | 200*3=600 | 49 |
| *yw; L^2^*/CyO*; ey-*GAL4*/ey-*GAL4 *X UAS-C4 /S-T* | *L^2^*/+*; ey-*GAL4*/C4* | 200*3=600 | 58.2 |
| *yw; L^2^*/CyO*; ey-*GAL4*/ey-*GAL4 *X UAS-C5 /S-T* | *L^2^*/+*; ey-*GAL4*/C5* | 200*3=600 | 38.7 |
| *yw; L^2^*/CyO;UAS-*p35*/Tb X +/+; *ey-*GAL4*/ey-*GAL4 | *L^2^*/+*; ey-*GAL4*/UAS-p35* | 200*3=600 | 22.7 |
| *yw; UAS-wg/CyO; UAS-C4/UAS-C4 X +/+; ey-*GAL4*/ey-*GAL4 | wg/*+ ; UAS-C4/ey-*GAL4 | 200*3=600 | 16.8 |
| *yw; UAS-wg/CyO; UAS-C4/UAS-C4 X L^2^/CyO; ey-*GAL4*/ey-*GAL4 | wg/ *L^2^; UAS-C4/ey-*GAL4 | 200*3=600 | 21 |
| *yw; UAS-arm/CyO; UAS-C4/UAS-C4 X +/+; ey-*GAL4*/ey-*GAL4 | *arm/+ ; C4/ey-*GAL4 | 200*3=600 | 21 |
| *yw; UAS-arm/CyO;UAS-C4/UAS-C4 X L^2^/CyO;ey-*GAL4*/ ey-*GAL4 | *arm/ L^2^; C4/ey-*GAL4 | 200*3=600 | 22.8 |
| *yw; UAS-sgg/CyO; TM-3/Tb X L^2^/CyO; ey-*GAL4*/ey-*GAL4 | *sgg/ L^2^; TM-3/ey-*GAL4 | 200*3=600 | 27.3 |
| *yw; UAS-sgg/CyO; C4/C4 X L^2^/CyO; ey-*GAL4*/ey-*GAL4 | *sgg/ L^2^; C4/ey-*GAL4 | 200*3=600 | 71.5 |
| *yw; UAS-dTCF^DN^/CyO; TM-3/Tb X L^2^/CyO; ey-*GAL4*/ey-*GAL4 | *dTCF^DN^/ L^2^; TM-3/ey-*GAL4 | 200*3=600 | 21.8 |
| *yw; UAS-dTCF^DN^/CyO; UAS-C4/UAS-C4 X L^2^/CyO; ey-*GAL4*/ey-*GAL4 | *dTCF^DN^/L^2^; C4/ey-*GAL4 | 200*3=600 | 70.6 |
| *yw; UAS-Porc^RNAi^/CyO; TM-3/Tb X L^2^/CyO; ey-*GAL4*/ey-*GAL4 | *Porc^RNAi^/ L^2^; TM-3/ ey-*GAL4 | 200*3=600 | 20.8 |
| *yw; UAS-Porc^RNAi^/CyO; C4/C4 X L^2^/CyO; ey-*GAL4*/ey-*GAL4 | *Porc^RNAi^/ L^2^; C4/ ey-*GAL4 | 200*3=600 | 68.1 |

*S-T (SM6TM6b-Tb) is a linked balancer on chromosome II and Chromosome III

**Supplementary figure legends**

**Figure S1**

**Validation of newt genes misexpression in the *Drosophila*.**
